# Claudin-4 modulates autophagy via SLC1A5/LAT1 as a tolerance mechanism for genomic instability in ovarian cancer

**DOI:** 10.1101/2024.01.18.576263

**Authors:** Fabian R. Villagomez, Julie Lang, Patricia Webb, Margaret Neville, Elizabeth R. Woodruff, Benjamin G. Bitler

## Abstract

Genome instability is key for tumor heterogeneity and derives from defects in cell division and DNA damage repair. Tumors show tolerance for this characteristic, but its accumulation is regulated somehow to avoid catastrophic chromosomal alterations and cell death. Claudin-4 is upregulated and closely associated with genome instability and worse patient outcome in ovarian cancer. This protein is commonly described as a junctional protein participating in processes such as cell proliferation and DNA repair. However, its biological association with genomic instability is still poorly-understood. Here, we used CRISPRi and a claudin mimic peptide (CMP) to modulate the cladudin-4 expression and its function, respectively in *in-vitro* (high-grade serous carcinoma cells) and *in-vivo* (patient-derived xenograft in a humanized-mice model) systems. We found that claudin-4 promotes a protective cellular-mechanism that links cell-cell junctions to genome integrity. Disruption of this axis leads to irregular cellular connections and cell cycle that results in chromosomal alterations, a phenomenon associated with a novel functional link between claudin-4 and SLC1A5/LAT1 in regulating autophagy. Consequently, claudin-4’s disruption increased autophagy and associated with engulfment of cytoplasm-localized DNA. Furthermore, the claudin-4/SLC1A5/LAT1 biological axis correlates with decrease ovarian cancer patient survival and targeting claudin-4 *in-vivo* with CMP resulted in increased niraparib (PARPi) efficacy, correlating with increased tumoral infiltration of T CD8+ lymphocytes. Our results show that the upregulation of claudin-4 enables a mechanism that promotes tolerance to genomic instability and immune evasion in ovarian cancer; thus, suggesting the potential of claudin-4 as a translational target for enhancing ovarian cancer treatment.

## INTRODUCTION

Ovarian cancer is the overarching term for a collection of unique diseases that represent the leading cause of death among gynecologic cancers. High-grade serous carcinoma (HGSC) is the most common and deadliest of these malignancies. Patients with HGSC initially respond to standard of care (surgery debulking, carboplatin, paclitaxel) and maintenance treatments (e.g., PARP inhibitors, PARPi). However, most of them eventually develop therapy resistance; therefore, overcoming such a resistance is an urgent need. ^1,2^

Genome instability is a hallmark of cancer; consequently, there is tolerance for such a characteristic within the physiological state of tumors. This hallmark is one of the factors that contribute to tumor progression, intratumorally heterogeneity, and the development of chemoresistance. ^1,3–5^ It arises from alterations in cell cycle progression and DNA damage repair (DDR) mechanisms, resulting in an increased rate of mutations and generation of multiple chromosomal modifications, such as amplifications, translocations, and micronuclei formation. Tumors are composed of heterogeneous cell subpopulations (metapopulation) exhibiting varying degrees of alterations in the genome; nevertheless, these cells can restrict the accrual of those alterations to prevent severe chromosomal transformations. Thus, this indicates the existence of a threshold tolerance for genomic instability within the tumor, and exceeding this critical threshold renders the tumor cells susceptible to cell death. However, the cellular mechanisms regulating such a tolerance are not well-known. ^3,6–11^

Claudin-4 is not normally expressed in the epithelium of the fallopian tube (which is believed to be the origin of HGSC) but is upregulated in up to 70% of HGSC tumors, ^12^ and its upregulation is associated with development of therapy resistance and worse patient outcome. This protein is traditionally though to regulate cell-cell junctions, more specifically tight junctions (a type of cell connections that are localized at the apical region of epithelial and endothelial cells); therefore, its primary function has been related to regulation of permeability. ^13,14^ In spite of that, it is known that claudin-4 also participates in multiple processes other than permeability, including migration, mitosis, apoptosis, and DNA repair. ^12,15,16^ Recently, we reported that in mitosis claudin-4 is associated in a way with proper dynamics of one of the components of the cytoskeleton, tubulin.^17^ Likewise, we showed that claudin-4 participates in a mechanism of DNA repair, the nonhomologous end joining, NHEJ and found that the levels of claudin-4 expression correlate with frequency of genetic modifications as well as therapy resistance in ovarian tumor samples.^12^ However, the molecular mechanims by which claudin-4 regulates genomic instability in ovarian cancer are still poorly understood. Here, we generated *in-vitro* (claudin-4 overexpression [OVCAR8 cells which do not express claudin-4]; and claudin-4 downregulation [OVCA429 and OVCAR3 cells which do express claudin-4] in HGSC cells) and *in-vivo* (PDX-HIS mice; patient-derived xenograft-human immune system mice) systems to modulate the function of claudin-4. We found that claudin-4 drives a dual regulatory mechanism on cell-cell junctions and cell cycle progression, contributing to the threshol-tolerance of genomic instability by ovarian tumor cells. Furthermore, targeting claudin-4 *in vivo* was found to enhance the treatment efficacy of a FDA-approved PARPi, niraparib in inhibiting tumor progression and increasing intratumural infiltration of T CD8+ lymphocytes.

## RESULTS

### Claudin-4 controls the heterogeneity of HGSC cells through fine-tuned cell cycle progression and inhibition of chromosomal amplifications.

Tumor heterogeneity is associated with diverse expression programs and biological functions, a phenomenon also observed in tumor-derived cell lines. Among these biological functions, the cell cycle stands out. ^18,19^ ^20^ This cellular process is significantly controlled by nutrient availability and ^21^ is intimately related to chromosomal instability. ^22,23^ Claudin-4 has been reported to participate in the cell cycle, but its biological role is still poorly-understood. ^17^ To address in more detail the biological role of claudin-4 in the cell cycle and genomic instability, we modulated its expression in different HGSC cells (*Supplementary table 1*). For this purpose, we used claudin-4 downregulation (knockdown, KD) and overexpression (claudin-4 overexpression, C4) systems, along with a claudin mimic peptide (CMP) designed by our laboratory. The peptide targets a homologous and conserved sequence located in the short extracellular loop in claudin-4 (*Supplementary* Figure 1a*, b*), thereby interfering with its interaction with other proteins. ^16,24–26^

First, we analyzed cycle progression by flow cytometry using wild type (WT) HGSC cells stained with propidium iodide. Under common culture conditions, the phenotype we observed is that HGSC cells cycle among all phases (G0-G1, S, and G2-M). Particularly, over time (24h *vs* 48h) some tumor cells stayed longer in the G0-G1 phase, downregulating their transit to the S and G2-M phases. Thus, we verified that the cell cycle progresses heterogeneously within both similar (same cell line) and different (OVCAR8, OVCA429, and OVCAR3) HGSC cells, fluctuating over time. This resulted in some cells not actively preparing for division (G0-G1) or clearly in mitosis (G2-M) (**Figure 1a**), potentially due to decreased nutrient availability (*Supplementary* Figure 1c). Manipulation of claudin-4 expression modified such heterogeneity in HGSC cells. Overexpression of claudin-4 associated with significant reduction of cells present in the S-phase (**Figure 1b**). Downregulation of claudin-4 correlated with significantly more cells in the G2-M phase and fewer in the G0-G1 phases in OVCA429 cells (**Figure 1c**), with a similar trend observed in OVCAR3 cells (**Figure 1d**). This observation is consistent with the higher number of OVCAR3 cells quantified in the G2-M phase (synchronized using starvation at 6h), as reported previously due to claudin-4 downregulation. ^17^ Collectively, our results show that claudin-4 significantly influences the heterogeneity of the cell cycle progression in HGSC cells by supporting their transit through various phases. Within this cellular process, it appears that the expression of claudin-4 arrests some HGSC cells in the G0-G1 phase, as indicated by more cells in the G0-G1 phase at 24h and 48h (**Figure 1b**), resulting in a smaller number of cells transitioning to the S phase (**Figure 1b**). This phenotype is associated with the observation of a finely-tuned exit from the G2-M phase, coupled with decreased entry into the G0-G1 phase observed in OVCA429 cells during claudin-4 downregulation (**Figure 1c**), also mirroring the phenotype observed in OVCAR3 cells (**Figure 2d**, with fewer cells in the G0-G1 and more in the G2-M phases at 24h).

**Figure 1.**
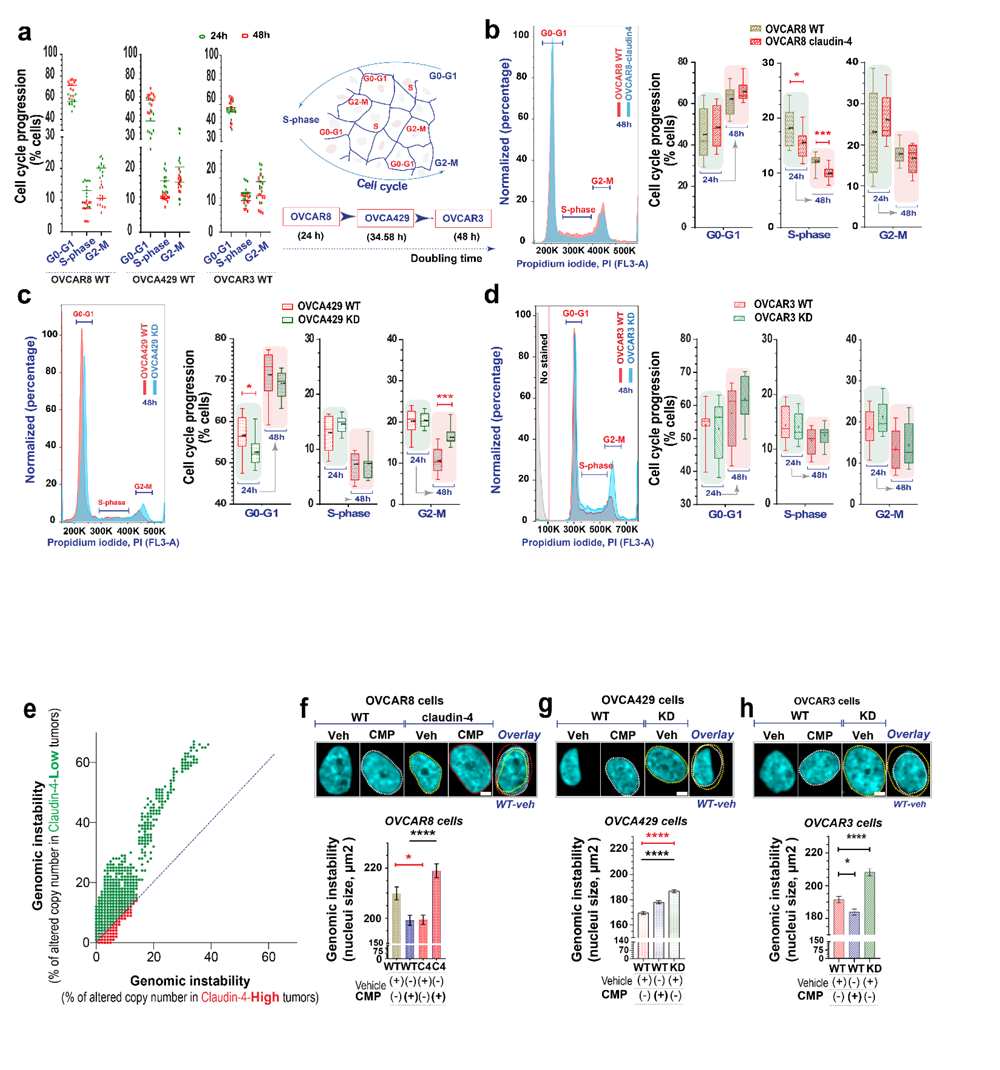
Claudin-4 accounts for intratumoral diversity through moderation of cell cycle and genomic instability in HGSC cells. Intratumoral diversity was explored by evaluating the transit of different HGSC cells through cell cycle. Cells were quantified and seeded without any stimuli followed by staining with propidium iodide (at 24h and 48h) and flow cytometry analysis. The association of claudin-4 with genomic instability was analyzed in human tumors (TCGA) and *in vitro* by treating cells with CMP (400µM) for 48h followed by single cell analysis of fixed (stained with dapi and phalloidin to mark genomic DNA). **(a)** Left, graphs show WT tumor cells in different phases of the cell cycle (green symbol at 24h; red symbol at 48h) which also changes as the time went by; right, drawing highlighting diversity of tumor cells withing the tumor, indicated as localization in different phases of cell cycle. Also, it highlights differences in doubling time among HGSC cells. **(b)** Left, representative histograms of cell cycle phases before claudin-4 over-expression; right, percentages of cells in every cell cycle progression. Similar data is indicated when claudin-4 expression was downregulated in OVCA429 **(c)** and OVCAR3 **(d)**. (4 independent experiments; Two-tailed Unpaired t test). **(e),** correlation of genomic instability (indicated as % of altered chromosomic copy numbers) in human ovarian tumors and claudin-4 expression. **(f),** top, confocal images showing (maximum projections) genomic instability (indicated as nuclei size, bottom: corresponding quantification) associated with claudin-4 overexpression or downregulation in OVCA429 **(g)** and OVCAR3 **(h)**. (n= OVCAR8, 1711 cells; OVCA429, 2630 cells; Two-tailed Mann Whitney test, Kruskal-Wallis test with Dunn’s multiple comparisons). (3 independent experiments; significance, p<0.05. Graphs show mean and SEM (standard error of the mean), scale bar 5µm.

**Figure 2.**
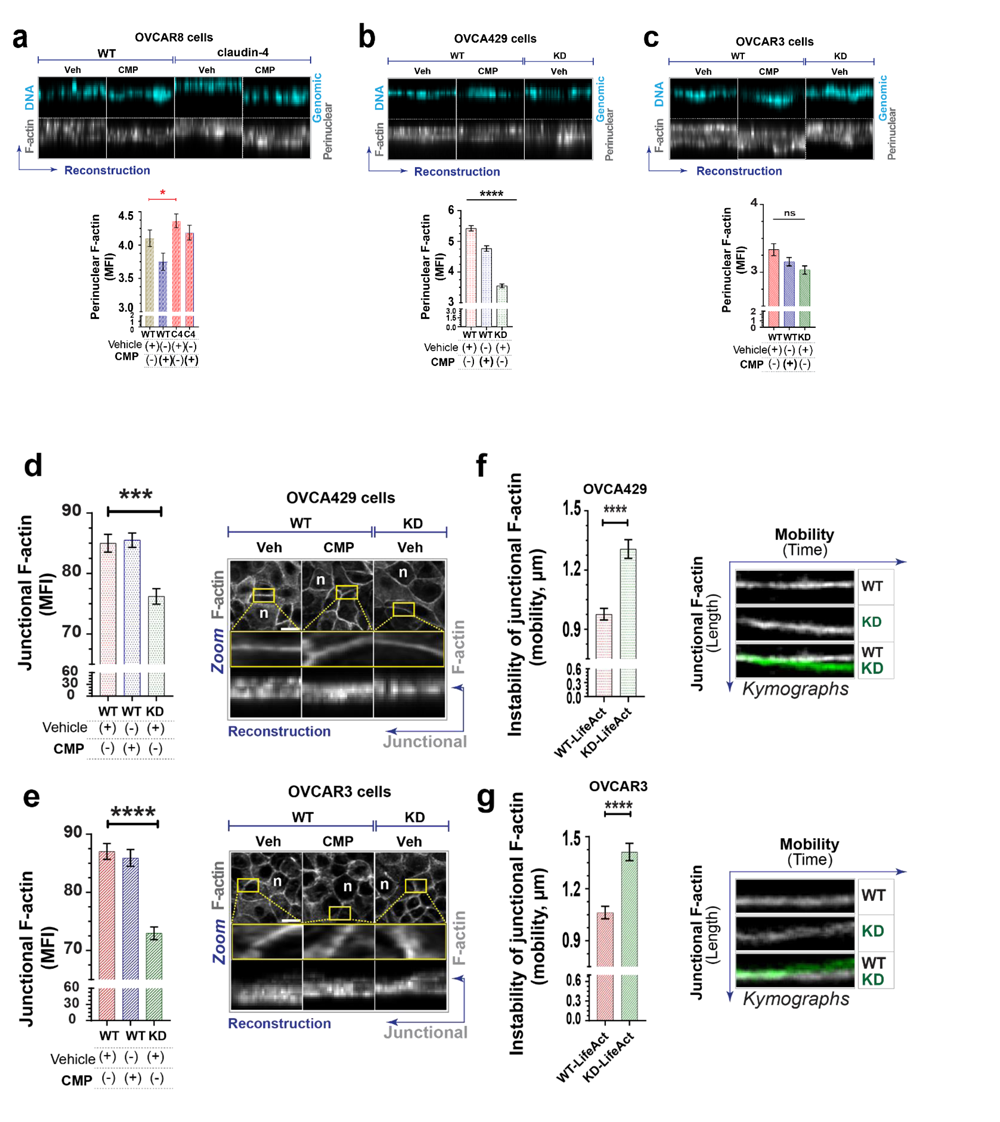
Claudin-4 stabilizes perinuclear and junctional F-actin in HGSC cells. The association of claudin-4 with the actin cytoskeleton was evaluated in fixed and living cells. HGSC cells were treated with CMP (400µM) for 48h and stained the actin-cytoskeleton with phalloidin. Also, HGSC cells were transduced to express LifeAtc (to mark actin in living cells). Then, we carried out a morphometric characterization. **(a)**, **(b)**, and **(c)** show reconstructions of perinuclear F-actin and genomic DNA (from confocal z-stacks) and corresponding quantification in OVCAR8, OVCA429, and OVCAR3 cells, respectively. (n= OVCAR8, 1711 cells; OVCA429, 2630 cells; OVCAR3, 2365 cells; Two-tailed Mann Whitney test, Kruskal-Wallis test with Dunn’s multiple comparisons). **(d)** and **(e)**, left, quantification of junctional F-actin from reconstructions (from confocal z-stacks); right, confocal images (maximum projection) and zoom, followed by reconstruction of selected regions of interest, ROIs (at junctional F-actin) from OVCA429 cells and OVCAR3 cells (bottom), respectively (n= OVCA429, 783 cells; OVCAR3, 825 cells; Kruskal-Wallis test with Dunn’s multiple comparisons). **(f),** kymographs generated from different ROIs (at junctional F-actin) from confocal live-cell imaging using transduced OVCA429 cells (n=142) with LifeAct to mark F-actin (without any stimuli and cultured for 24h) and OVCAR3 cells (bottom) (n=116) (Two-tailed Mann Whitney test). 3 independent experiments; significance, p<0.05. Graphs show mean and SEM (standard error of the mean), scale bar 5µm.

Additionally, it is known that an essential and shared characteristic between cell cycle progression and cancer development is nuclear morphological alteration. Cell cycle progression involves the remodeling of the nucleus as it advances through each phase, and its proper control is key to avoiding genomic instability. ^27–33^ Cancer development is related to significant changes in the morphology of the nucleus, which are intimately linked with genome instability. Particularly in ovarian cancer, there is a positive correlation between the nuclei size and chromosomic amplifications, a type of genomic instability. ^34–39^ Claudin-4 is associated with genomic instability ^12^, and in ovarian tumor samples, there is positive correlation between claudin-4 expression and reduced chromosomic amplifications (**Figure 1e**). Thus, indicating that claudin-4 somehow is limiting genomic instability. To gain further insights into this correlation, we used CMP to interfere with the function of claudin-4 and we examined indicators of genomic instability in HGSC cells through morphometric characterization of nuclei (stained with dapi). Targeting both claudin-4 expression and its function derived in significant morphological modifications in the nuclei of HGSC cells. The nuclear morphological architecture was reduced in claudin-4 overexpressing cells, and this phenotype was reversed by the effect CMP treatment, dependent on claudin-4 expression (**Figure 1f**). Conversely, this nuclear architecture was amplified when claudin-4 was downregulated in different HGSC cells (**Figure 1g, h**). However, the CMP effect on increasing nuclear size, was recapitulated only in OVCA429 cells. In contrast, the effect of CMP in OVCAR3 cells led to reduced nuclei size. This highlights not only the intratumoral diversity and a key biological role of the claudin-4 protein levels but also a significant effect of CMP in modifying genomic instability in HGSC cells, suggesting that the claudin-4-interacting proteins in the plasma membrane of OVCA429 and OVCAR3 could differ as well (*Supplementary* Figure 1b). Given that nuclei size is a signal of genomic instability, these initial results indicate that in our *in vitro* models, claudin-4’s expression exhibits a similar phenotype to that in HGSC tumor concerning genomic instability (**Figure 1e**). Additionally, these findings highlight the biological significance of claudin-4 in ovarian tumors, suggesting its possible regulatory role of proper cell cycle progression to prevent one of the indicators of genomic instability, chromosomic amplifications.

### Claudin-4 controls a physiological connection between genomic instability and the actin cytoskeleton in HGSC cells

The cytoskeleton is composed of elements such as actin, microtubules, and intermediate filaments, which are intimately related to cell-cell junctions in maintaining the morphology of epithelial and endothelial tissues. This structure physically connects nuclei with cell-cell junctions, specifically through junctional F-actin, and its dynamics are essential during cell cycle. ^27–29,31,33,40,41^ To further explore the regulatory role of claudin-4 in cell cycle and genomic instability, we stained HGSC cells with phalloidin to label the main component of the cytoskeleton, actin ^42^ and assessed its intracellular distribution, particularly at the perinuclear and cell junction regions. Our analysis revealed significant changes in the actin-cytoskeleton, characterized by an increased accumulation of perinuclear F-actin during claudin-4 overexpression (**Figure 2a**) and the opposite effect during its downregulation, notably in OVCA429 cells (**Figure 2b, c**). Additionally, all HGSC cells treated with CMP exhibited a trend of reduced F-actin localization at the perimeter of nuclei. This result implies that the proper functioning of claudin-4 is important for such localization. Further differences were observed at the cell junction region, where F-actin accumulated significantly less when claudin-4 was downregulated in various HGSC cells (**Figure 2d, e**). However, CMP treatment did not affect this accumulation, indicating that the adequate function of claudin-4 in maintaining junctional F-actin is less sensitivity to CMP inhibition compared to perinuclear F-actin, suggesting the relevance of claudin-4 in cell-to-cell connections.

Given that claudin-4 is traditionally described as a junctional protein, we hypothesize that the reduction of F-actin observed during claudin-4 downregulation in fixed cells could alter cell connections in living cells, thus changing tissue architecture and the phenotype of tumor cells, as reported. ^43,44^ Therefore, we generated HGSC cells expressing lifeact (a marker of F-actin) and performed time-lapse confocal imaging to analyze junctional F-actin dynamics as previously reported.^44^ As a result of claudin-4 downregulation, cell junctions moved more and irregularly over time. This observation is consistent with results observed in fixed cells (**Figure 2d, e**) and strongly suggest that claudin-4 downregulation makes cell-cell junctions more unstable and dynamic (**Figure 2f, g**; *Supplementary Figure1d*, *Supplementary video 1 and 2*). We conclude that the effect of claudin-4 in cell cycle and genomic instability is intimately related to its functional association with the actin-cytoskeleton, especially actin dynamics at the perinuclear and cell junction regions.

### The disruption of claudin-4 correlates with the formation of micronuclei and the activation of the cGAS-STING/autophagy molecular axis

The physical connection of nuclei with the cytoskeleton occurs through the linker of nucleoskeleton and cytoskeleton (LINC) complex, which binds to nuclear lamina and underlies the nuclear envelope. ^45^ Maintaining the integrity of the nuclear envelope (NE) is essential to avoid the release of nuclear content into the cytoplasm. However, this structure can be ruptured by pressure exerted by perinuclear F-actin, ^30,46,47^ which is enhanced when the function of p53 is lost. ^48^ These conditions can lead to the formation of micronuclei originating from different sources of DNA and phases of the cell cycle, such as interphase and mitosis. ^49–51^ Similar to the nucleus, micronuclei can also be affected by NE ruptures due to changes in lamin B1 and lamin A/C (the main component of nuclear lamina) dynamics. Consequently, DNA is more likely to be released into the cytoplasm, potentially activating the cGAS-STING pathway for an anti-tumor immune response. ^30,51–54^ We analyzed the possible association of claudin-4 with micronuclei formation and the cGAS-STING pathway in HGSC cells by confocal microscopy and immunoblotting.

We identified micronuclei in OVCAR8, OVCA429, and OVCAR3 cells, indicating that this type of chromosomic alteration is a common characteristic of HGSC cells. However, the expression and function of claudin-4 was found to be somehow related to micronuclei formation. Both overexpression and downregulation of claudin-4 correlated with significant changes in the frequency of micronuclei quantification. Specifically, a positive correlation was observed between claudin-4 overexpression and an increased number of micronuclei (**Figure 3a**), and a trend towards the opposite effect was noted during its downregulation in OVCA429 cells (**Figure 3b**). In OVCAR3 cells, a decrease in claudin-4 expression correlated positively with a higher incidence of micronuclei (**Figure 3c**). These differences could be attributed to the unique doubling time of OVCAR8 and OVCAR3 cells (**Figure 1a**) and to the similar trend we observed in the cell cycle between these cells (OVCAR8 and OVCAR3), both of which were more localized in the G2-M phase at 24h (**Figure 1b, d**). In addition, we quantified the total levels of lamin B1 and lamin A/C, finding that these proteins tended to vary before claudin-4 manipulation, particularly during claudin-4 downregulation (*Supplementary* Figure 1e-g). Furthermore, we characterized the nuclear envelope surrounding the micronuclei) in all HGSC cells and identified that some of these structures lacked components of the nuclear lamina (lamin B1, lamin A/C) (**Figure 3d-f**). Interestingly, during claudin-4 downregulation, the localization of lamin proteins in the nucleus changed significantly, notably with a higher amount of lamin B1 in HGSC claudin-4 knockdown cells (**Figure 3g, h**), indicating changes in lamin B1 dynamic. ^54^ This result correlates with the reduced perinuclear F-actin observed in KD cells (**Figure 2b, c**). Given that claudin-4 is localized in the nucleus in OVCAR3 ^55^ as well as in OVCA429 (not shown), this phenotype suggests that claudin-4 may modify nuclear lamina dynamics, possibly through a displacing mechanism. This mechanism is akin to the competition and exclusion observed between the actin bundling proteins, actinin and fascin, where they compete and exclude each other to bundle actin filaments into different actin networks. ^56^ Although the origin of micronuclei formation was not determined, our results show that claudin-4 impacts its formation or elimination, possibly through autophagy, as observed in osteosarcoma U2OS cells. ^57^ Moreover, the lack of nuclear lamina in micronuclei highlights the possibility that the DNA would be more likely to be released into the cytoplasm and activate the cGAS-STING pathway for an anti-tumor immune response. ^30,51,52^

**Figure 3.**
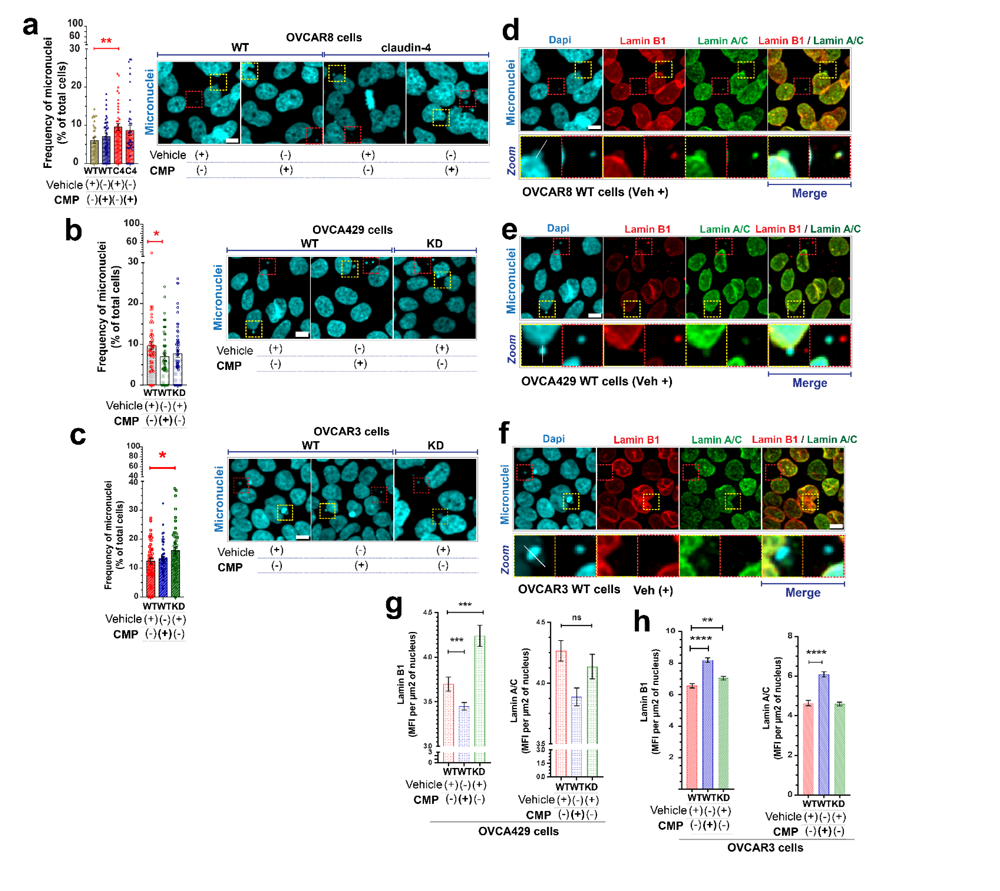
Claudin-4 modifies the structure of nuclear lamina. HGSC cells were treated with CMP (400µM; 48h) and fixed and stained with dapi, lamin B1 and lamin A/C. Characterization of nuclear lamina was carried out by confocal microscopy. **(a)** Left, frequency of micronuclei identification before claudin-4 over-expression; right, confocal images (maximum projections) highlighting (dotted squares) nuclei and micronuclei. Similar data is presented for claudin-4 downregulation in OVCA429 **(b)** and OVCAR3 cells **(c)**. (n= 4134 OVCAR8 cells; n=3315 OVCA429 cells; n= 4653 OVCAR3 cells; Two-tailed Mann Whitney test). **(d)** Selected confocal images (OVCAR8 WT cells treated with the vehicle from **(a)**) for the main components of nuclear lamina, lamin B1 and lamin A/C highlighting the lack of nuclear lamina in some micronuclei. Similar data is presented in **(e)** and **(f)** for OVCA429 and OVCAR3 cells, respectively. **(g)** and **(h)** Quantification of lamin B1 and lamin A/C in the nucleus in HGSC cells during claudin-4 downregulation. (two independent experiments; Kruskal-Wallis test with Dunn’s multiple comparisons, p<0.05). Graphs show mean and SEM, scale bar 5µm.

In ovarian cancer, the cGAS-STING pathway is typically inhibited as a mechanism of tumor immune evasion. ^52,58^ However, our data suggest that micronuclei in HGSC cells are likely to release DNA into the cytoplasm, activating the cGAS-STING pathway. Therefore, we evaluated the activation status of this pathway using immunoblotting and confocal microscopy. Briefly, upon cytoplasmic DNA detection by cGAS, cGAMP is produced, which then binds STING, recruiting TBK1. This process culminates in the phosphorylation TBK1 and STING (pTBK1 and pSTING), followed by downstream signaling of IRF3 to produce a type I interferon (IFN) response (**Figure 4a**). On the other hand, pSTING can induce autophagy through an independent mechanism of TBK1.^59,60^ Manipulating the function of claudin-4 resulted in changes in the phosphorylation status of STING and TBK1 (**Figure 4b**); suggesting that the activity of the cGAS-STING pathway was altered due to modifications in claudin-4 expression. The levels of pSTING were increased in both claudin-4 overexpression and downregulation, while the levels of pTBK1 (downstream of pSTING) were reduced under these conditions. This correlation, with more pSTING and less pTBK1, suggests that pSTING might be inactivated, leading to reduced pTBK1 or an increase in the type I IFN response. A recent report described that the levels of pSTING are regulated through direct encapsulation by lysosomes (micro autophagy). ^61^ Consequently, we stained HGSC cells for pSTING and lamp1 (lysosome marker) and visualized these cells by confocal microscopy. We found that pSTING was notably elevated before the claudin-4 downregulation (**Figure 4c**), consistent with our immunoblotting results. Moreover, in HGSC cells, we observed that pSTING was associated with lamp1, apparently inside lysosomes. This observation suggests that the direct encapsulation of pSTING by lysosomes could be occurring in HGSC cells as well. ^61^ This finding emphasizes the importance of the lysosome-mediated degradative pathway, as well as the necessity of vesicular intracellular trafficking in such a process, for the proper functioning of the cGAS-STING/Autophagy axis regulation. This notion is supported by our results in OVCA429 WT cells, which were manipulated to express ISRE (interferon-stimulated response element) to measure more directly the activation of a type I interferon IFN response. We treated these cells with cGAMP (an activator of cGAS-STING and type I IFN response) and found that this treatment increased a type I IFN response in OVCA429 cells; however, this induction was ablated when blocking vesicular trafficking using chloroquine (CQ) (*Supplementary* Figure 1h).

**Figure 4.**
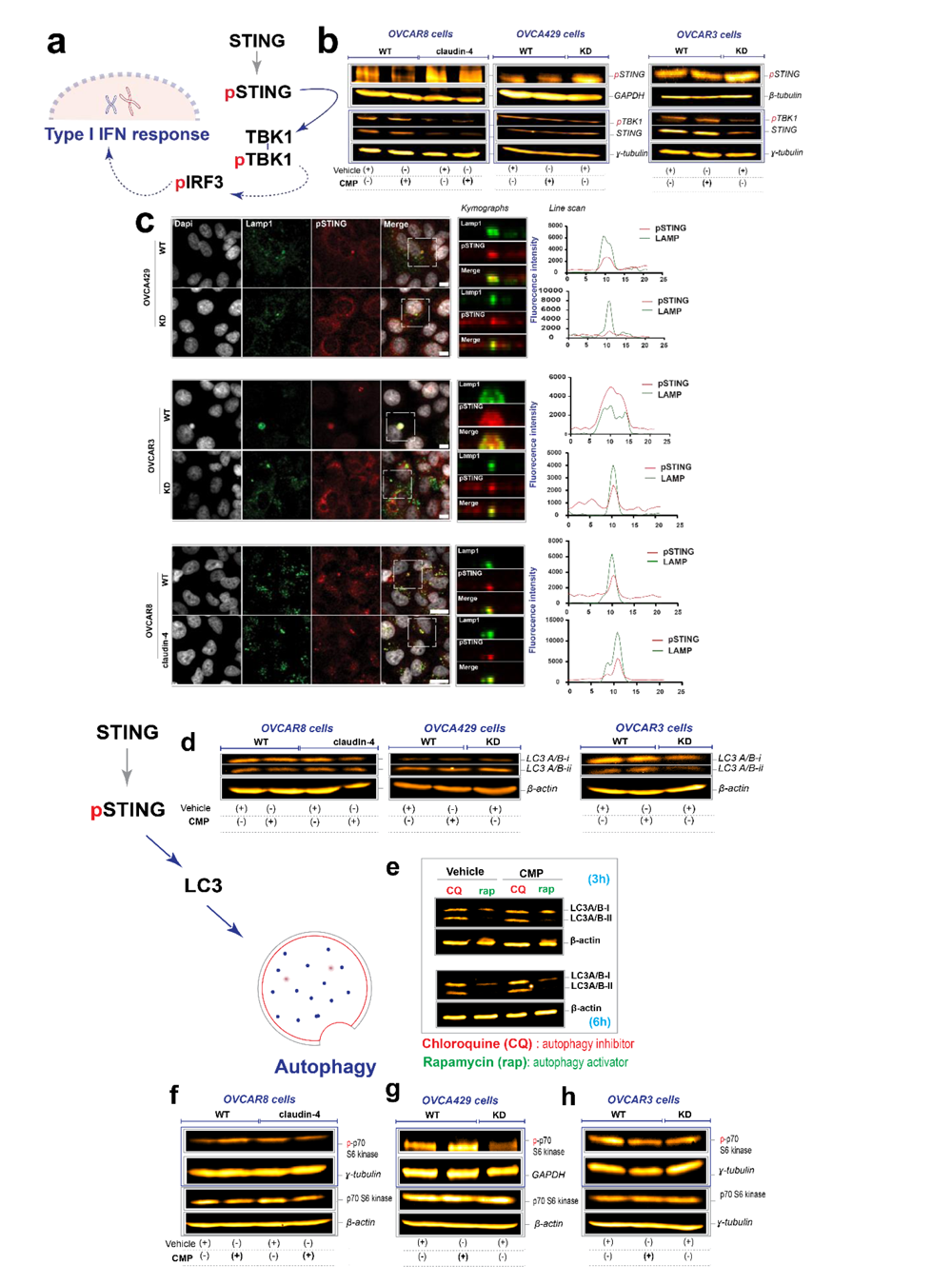
Claudin-4 impacts the cGAS-STING/autophagy molecular axis. Evaluation of the effect of claudin-4 on STING activation was performed via immunoblotting and confocal microscopy. Protein lysates were obtained from HGSC cells were treated with CMP (400µM; 48h) or stained for pSTING. **(a)** Drawing indicating the general signaling cascade of STING to activate a type I IFN response. **(b)** Immunoblotting for pSTING and pTBK1 in different HGSC cells. **(c)** Left, confocal microscopy of HGSC cells stained for pSTING and LAMP1; right, line scan from selected regions (dotted squares) showing similar patter of intracellular distribution of lamp1 and pSTING. **(d)** Left, drawing highlighting the association of STING with autophagy (LC3 marker of autophagy), independent of pTBK1; right, immunoblotting for LC3 in different HGSC cells. **(e)** immunoblotting for LC3 using lysates of OVCA429 cells treated with inhibitor (CQ, 40µM) or activator (rapamycin, 8µM) of autophagy. **(f)**, **(g)**, and **(h)** are immunoblotting for a downstream target of mTORC1, p70 S6 kinase (phospho-p70 S6 kinase) in different HGCS cells. Scale bar 10µm.

On the other hand, we performed immunoblotting for the autophagy marker, LC3 A/B, which has two isoforms: LC3 A/B-i (cytosolic) and LC3 A/B-ii (membrane bound). The lipidated LC3 A/B-ii form binds to autophagosomes, which are subsequently degraded as autophagosomes trafficked intracellularly and fuse with lysosomes. Our immunoblotting indicated that the manipulation of claudin-4 expression also affected autophagy (**Figure 4d**), seemingly by increasing this pathway. This notion is supported by the observation that when OVCA429 cells were treated with chloroquine (an autophagy inhibitor via blocking vesicular trafficking) or rapamycin (an autophagy activator via inhibition of mTORC1), we observed changes in the levels of LC3 A/B-ii, like those observed when autophagy was induced with rapamycin. (**Figure 4e**) Additionally, the phosphorylation status of a downstream target of mTORC1, p70 S6 kinase, changed due to claudin-4 manipulation (overexpression and downregulation) (**Figure 4f-h**). Specifically, these levels of phosphorylation were reduced in claudin-4 knockdown cells (**Figure 4g, h**), suggesting that the function of mTORC1 was inhibited. Together, our results demonstrate that claudin-4 significantly impacts the cGAS-STING/autophagy molecular axis, particularly suggesting its impact on the fate of micronuclei formation in HGSC cells.

### Micronuclei are intimately related to autophagy which is controlled via a functional connection of claudin-4 with the transporters of amino acids, SLC1A5/LAT1 in HGSC cells

The axis of cGAS-STING/autophagy has been associated with the clearance of cytosolic DNA,^60^ and removal of micronuclei has been indicated in osteosarcoma U2OS cells. ^57^ Moreover, claudin-4 appears to modulate micronuclei formation in some way. To confirm our hypothesis regarding the function of claudin-4 in micronuclei modulation through autophagy, we generated HGSC cells expressing mCherry-GFP-LC3 to measure autophagy activity (autophagy flux) using flow cytometry and confocal microscopy, as previously reported. ^62^ Briefly, LC3 serves as a marker of autophagy, localizing in vacuoles named autophagosomes. These vacuoles traffic intracellularly, fusing with lysosomes, causing a decrease in the intraluminal vacuolar pH and subsequently quenching of GFP fluorescence, unlike mCherry, that its resistance to such pH changes. Consequently, autophagy flux results in an increase in mCherry positive cells derived from double positive GFP-mCherry cells (*Supplementary* Figure 2a). To validate our strategy, HGSC-mCherry-GFP-LC3 cells were treated with CQ (chloroquine) and rapamycin to block and activate autophagy, respectively. These cells responded appropriately to these stimuli (*Supplementary* Figure 2b). Similarly, we treated our cells with CMP and analyzed autophagy using flow cytometry. We confirmed that claudin-4 participates in autophagy in HGSC cells. Both claudin-4 downregulation and overexpression correlated with increase autophagy activity (**Figure 5a-c**). This suggests that claudin-4 does not directly favor or limit the autophagy pathway but rather modulates it through other elements, e.g., mTORC1, the signaling complex regulating this activity, as suggested by our findings (**Figure 4f-h**). Remarkably, we found that micronuclei are engulfed through autophagy in HGSC cells (**Figure 5d-f**). Consequently, the possibility of DNA being released into the cytoplasm due to collapsed micronuclei is reduced, potentially representing a source to control genetic instability and nutrient supply. ^63^ Additionally, we found a similar correlation of autophagy and micronuclei *in vivo*. The marker of this process, LC3 A/B was closely associated to micronuclei (**Figure 5g**) and targeting claudin-4 with CMP showed signals of modification for that cellular pathway (**Figure 5h-i**, *Supplementary* Figure 3a-d). Our findings indicate a strong association of autophagy and the claudin-4’s functional association with genome instability.

**Figure 5.**
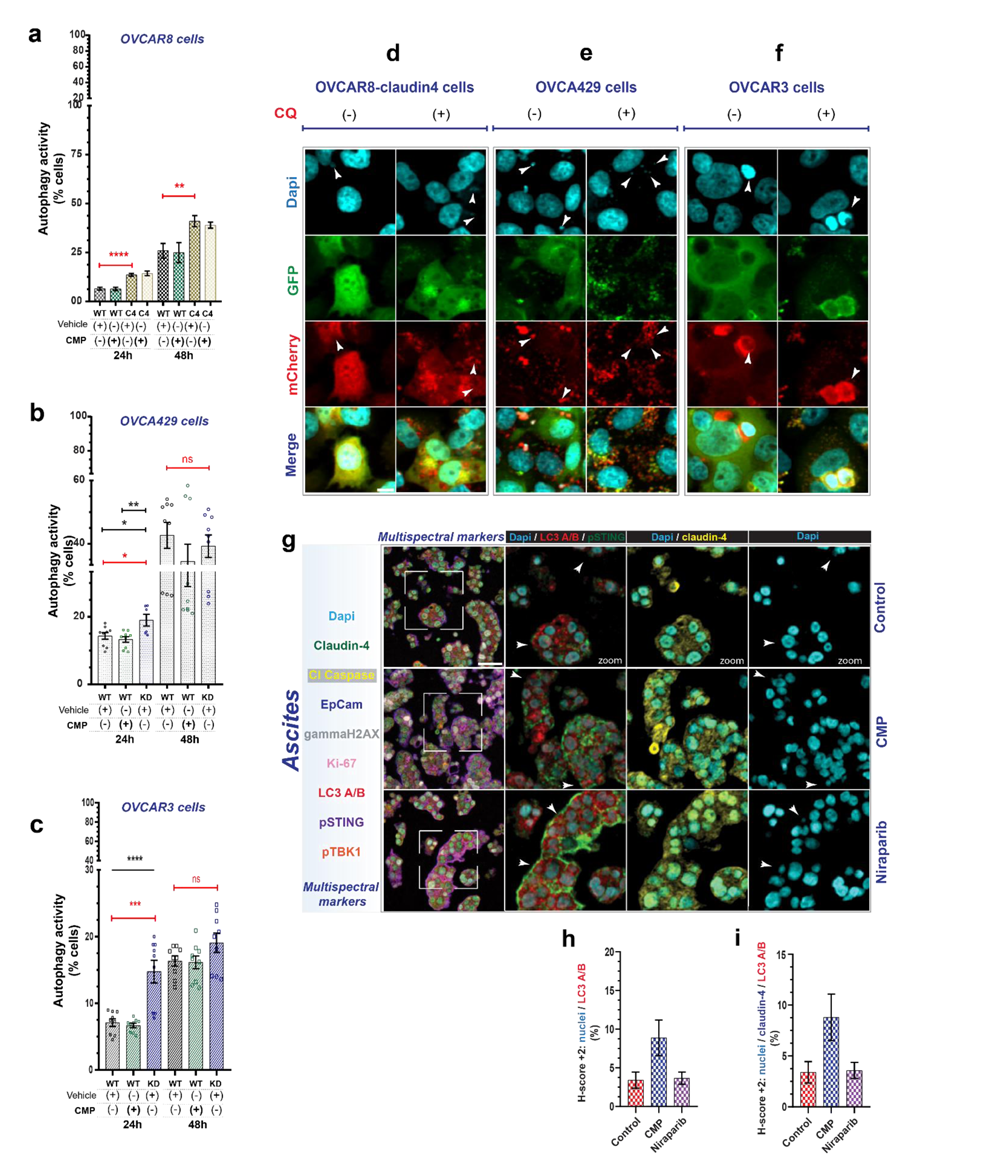
Claudin-4 modulates autophagy which is associated with micronuclei in HGSC cell lines and HGSC tumors. We analyzed the functional role of claudin-4 to link autophagy and micronuclei in HGSC cells, *in vitro* (HGSC-GFP-mCherry-LC3 expressing cells) and *in vivo* (patient derived xenograft in a humanized mice model) by flow cytometry, confocal microscopy, and multispectral immunofluorescence, IF. **(a)**, **(b)**, and **(c)** show percentages of HGSC cells with autophagy flux (by flow cytometry) before CMP treatment (400µM) and claudin-4 genetic manipulation (4 independent experiments). **(d)**, **(e)**, and **(f)** are confocal images (maximum projections) of HGSC-GFP-mCherry-LC3 expressing cells and stained with dapi (white arrow indicates the association of autophagy flux with micronuclei. **(g)** Ascites was obtained from tumor-bearing mice treated or not with CMP and prepared in histogel, followed paraffin embedding and multispectral staining. Representative multispectral IF images (all markers used are indicated in the blue-light box) are shown, where a region of interest (white square) is amplified to show specific markers (white arrow are highlighting the overlay of nuclei and the marker of autophagy, LC3 A/B). **(h)** and **(i)** indicate selected (+2) H-score quantification corresponding to percentage of positive nuclei with LC3 A/B, and nuclei with claudin-4 and LC3 A/B, respectively. (significance, p<0.05). Graphs show mean and SEM, scale bar 10µm and 50µm.

Afterwards, we focused on examining the molecular mechanism governing the claudin-4 role in autophagy. Previously, we identified the claudin-4-interacting proteins in OVCAR3 cells using BioID. ^55^ Some of the identified elements (as claudin-4-interacting proteins) are the transporters of amino acids SLC1A5, LAT1 (also known as SLC7A5), and SLC3A2. ^55^ The BioID method strongly suggests that claudin-4 and those proteins are in the same protein complex and possibly have a functional relationship. ^64^ It is known that LAT1 forms a heterodimer with SLC3A2 to enable a bidirectional transport system of amino acids along with SLC1A5. This system regulates upstream mTOR in cellular processes such as cell growth and autophagy which is mediated by a key role of L-glutamine influx/efflux.^65^

We employed different methodologies to assess the effect of claudin-4 function on the amino acid transport system and its association with autophagy regulation. Initially, we stained HGSC cells to visualize SLC1A5, LAT1, and claudin-4. By confocal microscopy, we determined that claudin-4 co-localized with both transporters of amino acids (**Figure 6a, b**). Additionally, we analyzed the protein expression of SLC1A5 and LAT by immunoblotting and found changes in SLC1A5 expression, especially in claudin-4 overexpression (*Supplementary* Figure 3e). However, the changes in LAT1 expression were significantly different derived from claudin-4 downregulation. There was a trend of less and more LAT1 expression in OVCAR8 claudin-4 cells (claudin-4 overexpression) and in OVCAR3 KD cells (claudin-4 downregulation), respectively. The opposite effect (more expression) was determined in OVCA429 KD cells (**Figure 6c**, *Supplementary* Figure 4a). This highlights differences among HGSC tumor cells as well as the relevance of claudin-4 association with the bidirectional transport system formed by SLC1A5/LAT1 (**Figure 6d**), with a particular association with LAT1 in such tumor cells. Furthermore, for the activity of the transport system of amino acids constituted by SLC1A5/LAT1, a key regulatory element is L-glutamine, which is internalized by SLC1A5 to increase its intracellular concentration. The amino acid then functions as a substrate for LAT1, which internalizes essential amino acids as L-glutamine is exiting the cell. Consequently, we limit L-glutamine availability in culture media and measure autophagy flux. All our HGSC cells exhibited increased autophagy flux due to L-glutamine limitation (**Figure 6e**) which agrees with previous reports. ^65^ Likewise, we observed by confocal microscopy changes in SLC1A5 intracellular distribution directly associated to claudin-4 overexpression and downregulation. The most significant difference was identified in OVCA429 KD cells, where SLC1A5 was markedly localized at the level of cell-cell junctions **(Figure 6f)** and interestingly, L-glutamine limitation was associated with changes in the SLC1A5 intracellular distribution, particularly in OVCA429 WT cell. In these cells, SLC1A5 localized clearly at the level of cell junctions because of L-glutamine limitation (**Figure 6g**). Additionally, because previously we observed accumulation of pSTING, which is involved in autophagy, in claudin-4 KD cells we cultured HGSC cells in media without L-glutamine and stained for pSTING and lamp1, as previously indicated (*Supplementary* Figure 4b). L-glutamine limitation correlated with accumulation of pSTING in OVCA429 and OVCAR3 WT cells, similarly to KD cells cultured in media with L-glutamine (**Figure 4c**). Also, autophagy blocking with CQ or autophagy induction with rapamycin was associated with changes in the intracellular distribution of SLC1A5 (*Supplementary* Figure 4c*, d*).

**Figure 6.**
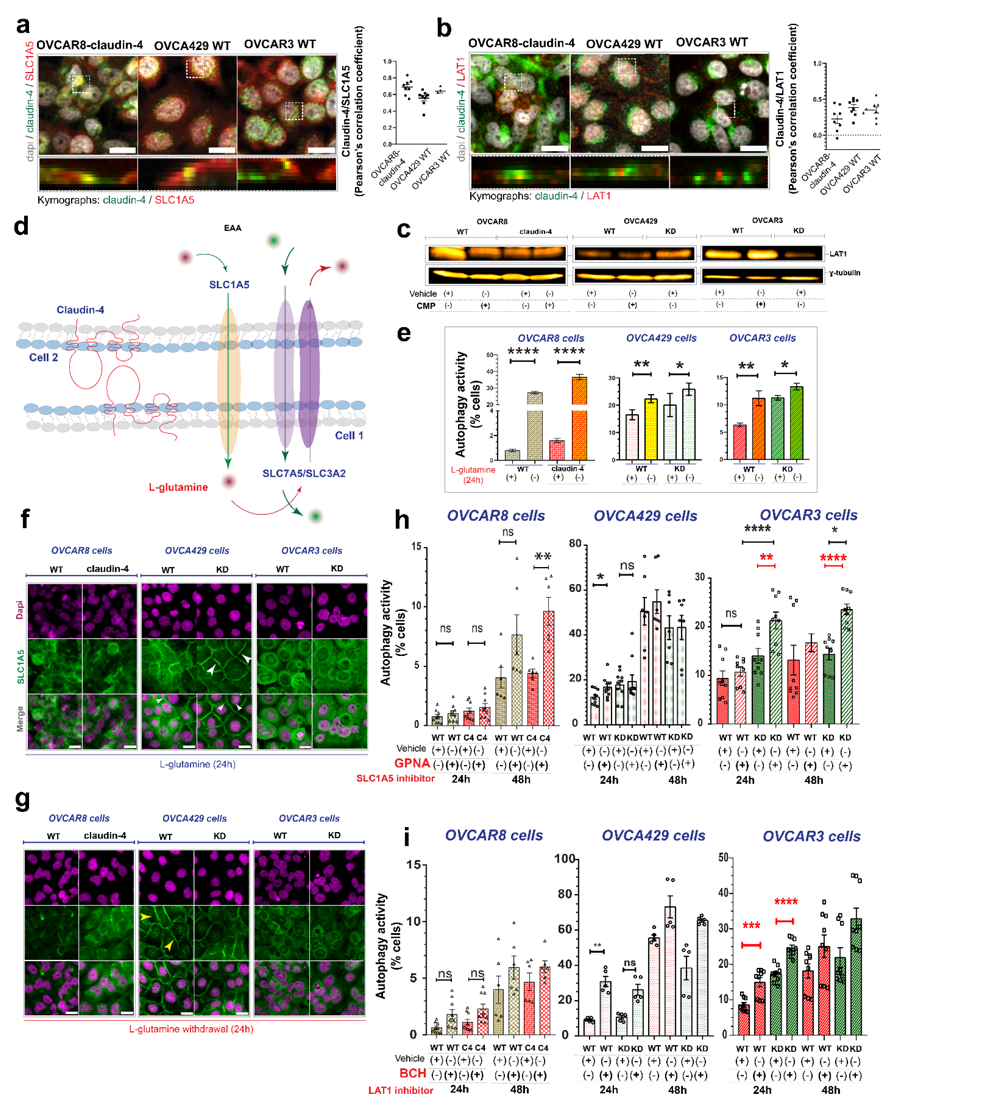
Claudin-4 forms a regulatory axis with SLC1A5/LAT1 to modulate autophagy in HGSC tumor cells. We studied the reported regulation of autophagy by SLC1A5/LAT1 and its association with claudin-4 in different HGSC cells. Modulation of claudin-4 function was associated with the activity of SLC1A5 and LAT1 using specific inhibitors and by limiting L-glutamine availability (a key regulator of the regulatory activity of SLC1A5/LAT1 on autophagy) all of which was studied by immunoblotting, confocal microscopy, and flow cytometry. **(a)** left, confocal images (maximum projections) showing the intracellular distribution of SLC1A5 and claudin-4 in HGSC cells; right, colocalization of SLC1A5 and claudin-4. **(b)** left, confocal images (maximum projections) showing the intracellular distribution of LAT1 and claudin-4 in HGSC cells; right, colocalization of LAT1 and claudin-4. **(c)** Shows representative immunoblots for LAT1 in HGSC cells before claudin-4 disruption (4 independent experiments). **(d)** Model of a functional association of claudin-4 with the bidirectional transport system formed by SLC1A5/LAT1 (adapted from ^65^) which propose that such association is important for proper transport of amino acids in HGSC cells. **(e)** Percentage of HGSC cells with autophagic flux before L-glutamine withdrawal and measured by flow cytometry. **(f)** and **(g)** show representative confocal images (maximum projections) of the intracellular distribution of SLC1A5 before claudin-4 overexpression (OVCAR8 claudin-4) and downregulation (OVCA429 KD and OVCAR3 KD) in HGSC cells before L-glutamine withdrawal. **(h)** Graphs indicate percentages of cells with autophagy flux when treated with specific inhibitor of SLC1A5 activity (GNPA, 5mM). **(i)** Graphs indicate percentages of cells with autophagy flux when treated with specific inhibitor of LAT1 activity (BCH, 25mM). (Two-tailed Unpaired t test and Mann Whitney test; Kruskal-Wallis test with Dunn’s multiple comparisons; One way ANOVA and Tukey’s multiple comparison test; 3 independent experiments; significance, p<0.05). Graphs show mean and SEM, scale bar 20µm.

Finally, to verify specificity, we inhibited the transporter function of SLC1A5/LAT1 as previously described, ^65^ using L-g-glutamyl-p-nitroanilide (GPNA) and 2-aminobicyclo-(2,2,1) heptane-carboxylic acid (BCH), respectively and analyzed such inhibition on autophagy activity. Inhibition of SLC1A5 and LAT1 (**Figure 6h, i**) resulted in a trend of increase autophagy flux in HGSC cells, particularly in OVCAR3 cells; thus, confirming that specific inhibition of SLC1A5 and LAT1 dysregulates autophagy in HGSC cells. Our data strongly suggests that the bidirectional transport system mentioned before is present in HGSC cells and that claudin-4 is functionally associated with it. For instance, by stabilizing cell-cell junctions when that system is formed at that location, as suggested in **Figure 6d**.

### Targeting claudin-4 *in vivo* modifies the tumor immune microenvironment and increases the therapy efficacy of niraparib in PDX-HIS mice

Previously, we targeted claudin-4 *in vivo* with CMP using immunodeficient tumor-bearing mice. Disruption of claudin-4 resulted in reduction of tumor growth due to cell death.^16^ However, the immunological consequences of targeting claudin-4 in tumors, via CMP are not known. To study the outcome of such intervention, we generated a humanized mouse model, the human immune system (HIS)-mice as previously reported.^66^ This model is considered a better translational approach than immunocompromised mice because it incorporates the contribution of a human immune system in modeling the human tumor progression. ^67^ We implanted a reported patient-derived xenograft (PDX, GTFB1016)^68^ into the humanized mice to get our *in vivo* (PDX-HIS-mice) system. After tumor establishment (**Figure 7a**, right; IVIS scan, before treatment), we applied cycles of treatments to the tumor-bearing mice with CMP (<u>claudin mimic</u> <u>peptide</u>) and follow-up tumor progression by IVIS scan. This this intervention was compared with a PARPi therapy, niraparib (<u>FDA-approved</u>) and with a combinatory therapy, CMP/niraparib (see methods and *Supplementary* Figure 6a*, b*).

**Figure 7.**
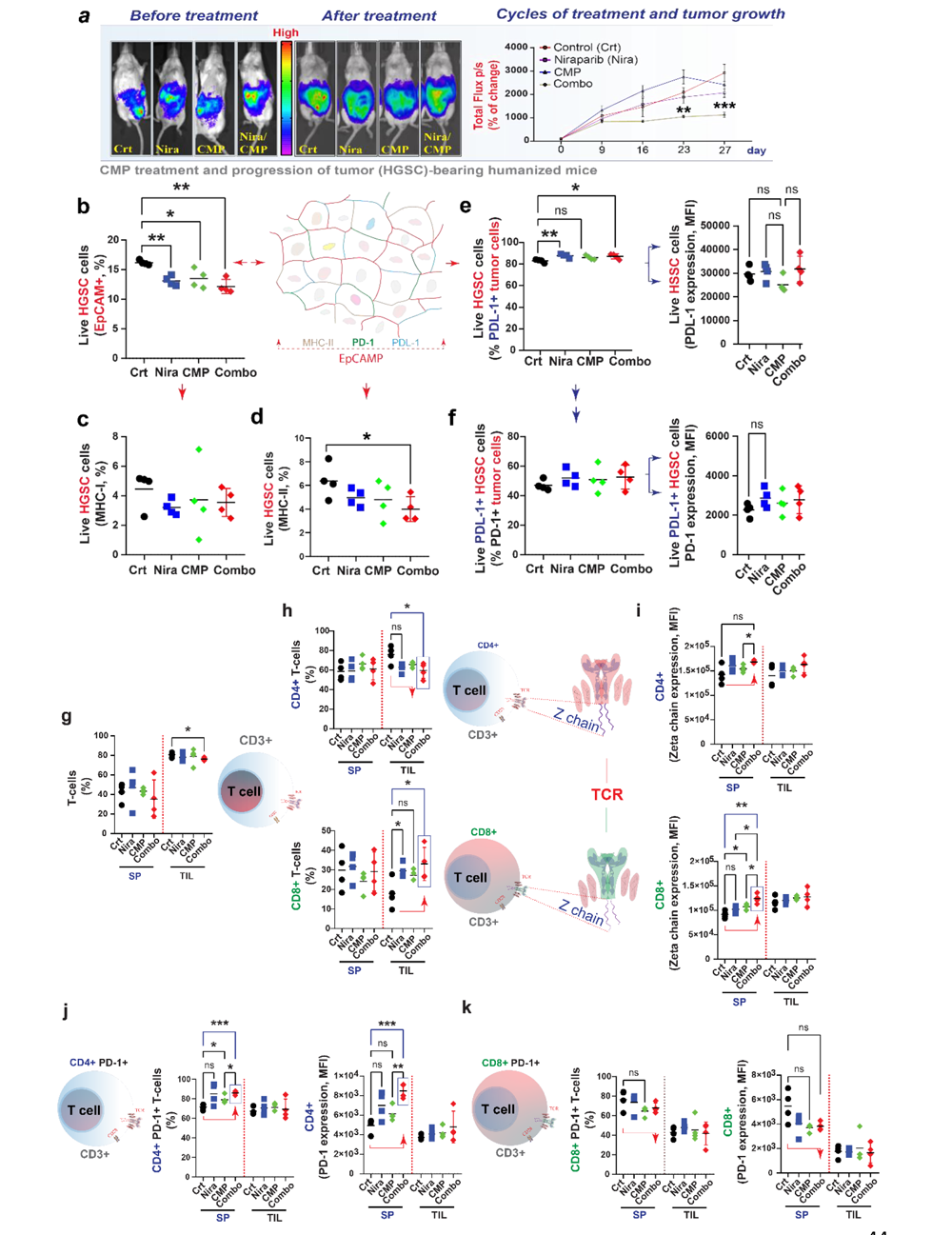
Targeting claudin-4 *in vivo* with CMP increases tumor-infiltrating T CD8+ lymphocytes and improves the therapy efficacy of niraparib in HGSC. We generated humanized mice and implanted a patient derived xenograft. After tumor establishment, mice were treated with CMP (4mg/kg), niraparib (Nira, 50mg/kg), and combination of CMP and niraparib (combo, 4mg/kg; 50mg/kg, respectively). At the end of treatment mice were sacrificed and we analyzed tumors and immune cells by flow cytometry (from *in vivo* experiment #2). **(a)** left, verification of tumor establishment by IVIS scan (before treatment); center, comparison of tumor size before treatment (day 0) *vs* after treatment (day 27); right, follow-up of tumor progression during therapy (indicated as bioluminescence signal over time; red color indicates high tumor growth in the IVIS scan). **(b)** left, live tumor cells recover after treatments; right, drawing highlighting that EpCAM protein was used as a tumor marker and red arrows indicate a gate from this marker. Also, PD-1, PDL-1 and MHC-ii with different color indicate that there are clones of tumors cells expressing these proteins **(c)** and **(d)** show percentage of live tumor cells positive for MHC-I and II, respectively. **(e)** left, percentage of live PDL-1+ tumor cells; right, corresponding protein expression by MFI (median fluorescence intensity). **(f)** left, percentage of live PDL-1+ tumor cells with PDL-1 expression (PDL-1+/PDL-+); right, corresponding PD1 expression by MFI. **(g)** Prevalence of T cells (CD3+) in tumor and spleen. **(h)** top, frequency of T CD4+ cells; bottom, frequency of T CD8+ cells. **(i)** top, z chain expression (MFI) on T CD4+ cells; bottom, z chain expression (MFI) on T CD8+ cells. **(j)** left, quantification of T CD4+ PD-1+ cells; and right, PD-1 expression (MFI). **(k)** left, numbers of T CD8+ PD-1+ cells; and right, PD-1 expression (MFI). Graphs show mean (every symbol represents data from a different mouse) (n=Control, 4 mice; CMP, 4 mice; niraparib, 4 mice; niraparib/CMP, 5 mice; Brown-Forsythe ANOVA test and Unpaired t with Welch’s correction; p<0.05).

As expected from our previous study, cycles of CMP treatment decreased tumor growth, ^16^ similar to cycles of niraparib treatment (*Supplementary* Figure 3f). Remarkably, combinatorial therapy showed a significant improvement in the effectiveness of niraparib, thus suggesting that claudin-4 function is associated with resistance to niraparib treatment *in vivo* (**Figure 7a**). Likewise, the physiological state of tumors (detected by EpCAM expression) was modified due to the therapeutic intervention, particularly in the CMP/niraparib combination. We quantified lower percentages of live tumor cells that were recovered from ascites samples (**Figure 7b**). As the same methodology was applied to process all samples, these results suggest that CMP and niraparib treatments somehow affected tumor survival, possibly by increasing tumor cell apoptosis. ^16,69^ In addition, we identified that the frequency of cells expressing MHC-I (major compatibility complex class-I) (**Figure 7c**) within the tumors was similar among all conditions, but not MHC-II (major histocompatibility complex class-II) which was not only expressed by tumors (MHC-II is usually expressed in professional antigen presenting cells, such as dendritic cells and B-cells which stimulate the activation of T CD4+ cells), ^70^ but its localization was significantly decreased in tumor cells treated with CMP/niraparib (**Figure 7d**). In different types of cancer, including ovarian cancer, the expression of MHC-II is associated with a better patient outcome, highlighting (as in other cancer types) the relevance of the T CD4+ lymphocyte activity as well as T CD8+ cells. ^70–75^

Furthermore, PDL-1+ tumor cells were observed more frequently in all groups of treated mice, especially in niraparib and CMP/niraparib treated tumor-bearing mice, with the particularity that the expression level of this protein slightly decreased in the CMP group **(Figure 7e)**. Additionally, these tumor cells expressed MHC-II with a similar trend to that observed in total live HGSC cells (*Supplementary* Figure 6a). The expression of this protein has been reported previously in ovarian cancer and related to a population of tumor cells with a stem cell phenotype. ^76^ This protein (PDL-1) is traditionally associated with suppression of immunity by tumor cells, as a cellular mechanism of tumor immune evasion, where PDL-1 can interact with its receptor (PD-1), which is upregulated in T cells when they are activated and functions a mechanism of immune cell regulation. ^77^ For example, it has been reported that high expression of PDL-1 in tumor cells and its interaction with PD-1 in T cells, interfere with signaling cascade (zeta chain of the TCR, T cell receptor and CD28) necessary to sustain the lymphocyte activation. ^78^ Strikingly, regardless of the treatment applied, tumors with PDL-1 expression co-expressed PD-1 as well, approximately 50%, **(Figure 7f)**. The co-expression of PDL-1 and PD-1 has been previously reported in tumors, where such co-expression is associated with a mechanism of tumor suppression associated to modulation of mTOR signaling and independent of adaptive immunity. ^79–82^ Together, our results show that the combination of CMP and niraparib treatment, modifies the physiological state of tumors by regulating their cellular composition; particularly limiting the increasing of tumor cells with MHC-II and increasing the number of tumor cells with PDL-1 and PD-1 co-expression. Thus, affecting the known systemic immunosuppression induced in cancer ^83^.

During immune surveillance, immune cells migrate through different lymphoid organs, notably the spleen ^84,85^. It is known that in this organ, T CD4 lymphocytes cooperate with T CD8 lymphocytes to support their activation ^86^ and enhancement of tumor destruction ^87^.

However, the spleen also represents a site of tolerance to cancer development which is importantly associated to the function of some myeloid cells. ^88,89^ Interestingly, derived from our treatments we observed reduced proportions of myeloid cells in the spleen, particularly when combining CMP and niraparib (*Supplementary* Figure 7b). Other immune cells such as B cells and natural killer cells were quantified as well (*Supplementary* Figure 7c and d*, respectively*). Specifically in T cells (**Figure 7g**), we observed increased infiltration of T CD8+ lymphocyte in tumor-bearing mice treated with CMP and this infiltration was significantly augmented using the CMP/niraparib combination (**Figure 7h**). This correlates with decreased tumor progression and the beneficial effect of tumoral infiltration of T CD8+ lymphocyte. ^90^ Likewise, the expression level of the zeta chain of the TCR (T cell receptor) was increased in lymphocytes (**Figure 7i**). This chain is usually decreased in different types of cancer, ^91^ including ovarian cancer. ^92^ This decrease is associated with reduced activity of lymphocytes and therefore, is considered a mechanism of tumor immune evasion. In the spleen a T CD4+ population of lymphocytes expressing PD-1 was significantly increased (**Figure 7j**) meanwhile the T CD8+ lymphocytes expressing PD-1 showed a trend to be reduced (**Figure 7k**). This suggests that these lymphocytes, at least in the spleen, the activation status of these cells is different, where it seems that T CD4+ cells are more active. Also, to mount a proper immune response, T CD8+ lymphocytes need the help of T CD4+ cells. ^92^ Then, is feasibly that the expression of the zeta chain of the TCR (in lymphocytes of the spleen) is a key element contributing to the help provided by T CD4+ to T CD8+ lymphocytes.

Collectively, our results suggest that the combined CMP/niraparib treatments select specific population of tumor cells characterized by co-expression of PLD-1/PD-1 and expression of MHC-ii. These selected cells appear to be less efficient in maintaining a systemic immunosuppression state. Consequently, fewer myeloid cells accumulate in the spleen, which hinders their ability to inhibit the function of lymphocytes. Furthermore, these lymphocytes (localized in the spleen) express an increased level of zeta chain (of the TCR), making them less susceptible to myeloid cell inhibition.

In summary, the CMP/niraparib treatment seems to influence the selection of a distinct composition of tumor cells within the tumor, marked by co-expression of PDL-1 and PD-1, along with MHC-ii. This selected population demonstrated reduced efficacy in inducing systemic immune suppression, particularly evident at the level of spleen and myeloid infiltration. Moreover, these tumor cells show a diminish ability to inhibit the expression of the zeta chain of the TCR in lymphocytes, rendering them more susceptible to recognition by immune cells.

A notable consideration is that the reduced presence of myeloid cells in the spleen, associated with decreased tolerance to cancer, coincides with heightened zeta chain expression in lymphocytes withing this secondary lymphoid organ. In contrast, these lymphocytes, particularly T CD4+ lymphocytes are likely to cooperate more effectively with T CD8+ lymphocytes, facilitating their infiltration into the tumor (*Supplementary* Figure 6e).

### The claudin-4 expression and function are highly linked with LAT1, which is associated with transport of amino acids and decreased survival of patients with HGSC

Here we show that the claudin-4 association with genomic instability in HGSC is intimately related to its role in cell cycle progression and modulation of genomic instability through autophagy regulation mediated by the transporters of amino acids, SLC1A5/LAT1. Furthermore, targeting claudin-4 modifies the tumor microenvironment resulting in decreased tumor progression and tumor immune evasion. To explore the implications of our *in-vitro* and *in-vivo* findings in HGSC tumor, we looked for data mining. First, we construct the protein-protein network of the claudin-4-interacting proteins we reported previously in OVCAR3 cells. ^55^ From such a network, we selected the proteins whose interaction is experimentally reported and then constructed another protein-protein network to select proteins with interacting reported experimentally. Together, these selected proteins were analyzed in publicly available data sets of HGSC tumors (cBioPortal). We selected those proteins that show higher mutation rates, the top for highly mutated proteins were BRD4, CCDC130, PTK2, and NDRG1. Afterwards, we included these proteins with those from our BioID results (whose interaction with claudin-4 was reported experimentally) to generate clusters of protein-protein networks (as indicated above) and look for biological processes. The distinctively associated cellular processes with such clusters were regulation of cell-cell junctions and cytoskeleton dynamics, and transport of amino acids through the plasma membrane (**Figure 8a**). Thus, it is highly possible that those cellular mechanisms are key drivers of the correlation of claudin-4 upregulation and a worse patient outcome in HGSC. This notion is supported by our findings regarding the correlation of expression, of the elements forming every functional cluster with claudin-4, with aggressiveness of HGSC tumors and decrease patient survival (**Figure 8b**). Likewise, we used these clusters (all, cluster 4, and cluster 4) to explore if they were correlated with aggressiveness in other types of cancer. We found that all clusters are associated with worse patient outcomes in breast and lung cancer but not in stomach. Notably, cluster 4 correlated with decreased RFS (relapse-free survival) and OS (overall survival) in breast and lung cancer, respectively (**Figure 8c**).

**Figure 8.**
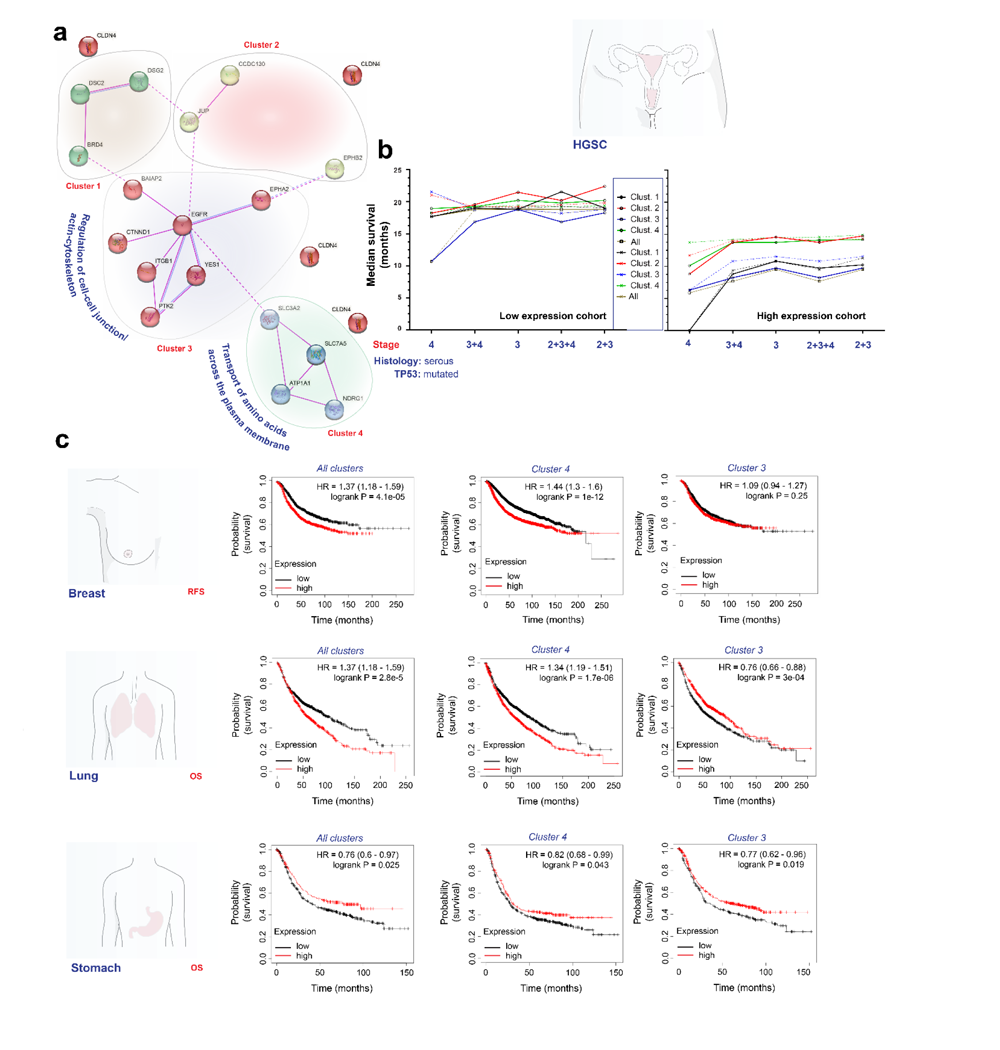
A protein cluster formed by claudin-4 associates with transport of amino acids and worse patient outcome in HGSC patient survival and possibly other cancer types. Reported claudin-4-interacting proteins were used to generate protein-protein interacting networks to find key elements and cellular functions associated with claudin-4 upregulation in HGSC tumors and publicly available data from HGSC tumor (cBioPortal and Kaplan-Meier plotter). **(a)** Protein-protein network (based on BioID) of claudin-4 interacting proteins (experimentally determined using STRING) and highly mutated proteins (cBioPortal) in HGSC tumors shows clusters of proteins and distinctive associated cellular functions. **(b)** Median survival of HGSC patients (criteria: p53 mutated) correlated with every cluster (based on clausin-4) identified (continue lines) and without claudin-4 (dotted lines) (p values and false discovery rate, FDR in indicated in *Supplementary table 2 and 3*). **(c)** Correlation analysis of relapse-free survival (RFS) in breast cancer and overall survival (OS) in lung and stomach cancer with identified claudin-4 clusters (all, cluster 4 and cluster 3).

## DISCUSSION

Genome instability is a hallmark of cancer, indicating that tumor cells have cellular mechanisms of tolerance to genomic instability. However, it is not well-know how tumor cells limit the accumulation of such modifications to avoid catastrophic chromosomic alterations and cell death. Here we show that claudin-4, traditionally described as a tight junctional protein, mediates a mechanism of tolerance to genomic instability in HGSC cells. This mechanism is a dual protective cellular mechanism that allows HGSC cells proper transit throughout the cell cycle phases and modulation of the accumulation of genomic instability via induction of autophagy.

Briefly, claudin-4’s biological role in cells includes the stabilization of the actin-cytoskeleton, particularly the F actin at the cell-cell junctions and perinuclear regions. All of these impacts the nuclear shape and cell cycle progression, and therefore, the modulation of genomic instability in HGSC cells.

Interestingly, micronuclei generated in HGSC cells are modulated by the function of claudin-4. These forms of genetic alterations are typically associated with the cGAS-STING activation, ^52^ but we show that they are also intimately related to autophagy. Consequently, in spite that micronuclei generated in HGSC cells can collapsed due to disruption of the nuclear envelope, the possibility of releasing the DNA contained these micronuclei into the cytoplasm is reduced because of autophagy induction, which can prevent micronuclei accumulation. ^93^

Furthermore, we identified a novel functional link of claudin-4 with the transporters of amino acids, SLC1A5 and LAT1 which regulate a bidirectional transport system of amino acids that control autophagy activation.^65^ Likewise, we found that such a functional link, particularly between claudin-4 and LAT1, strongly suggest that one key cellular process driving aggressiveness in ovarian cancer in the transport of amino acids through the plasma membrane. This is interestingly because formation of ascites in common in advanced stage of ovarian cancer and ascites play a fundamental role in immune regulation via amino acids and their transport ^94–96^

Remarkably, *in vivo* targeting of claudin-4 not only affected tumor progression but also such targeting, improved the efficacy of niraparib treatment for HGSC tumors. This effect was related to immune modulation of some tumor clones. For example, tumor cells with double expression of PD-1 and PDL-1 were selected due to treatment. Likewise, some cells from the HGSC tumor expressed MHC-11 but this was modulated differentially before disruption of claudin-4 and niraparib treatment. Thus, suggesting a that modulation of MHC-ii has a biological relevance for ovarian cancer progression. Also, we observed more infiltration of T CD8 lymphocytes in tumor-bearing mice treated with CMP/niraparib. This is consistent with the reported benefit of such infiltration in ovarian cancer. ^90^ Interestingly, we also identified that the zeta chain of the TCR (T cell receptor) was increased when tumor cells received treatment (CMP, niraparib, and combination), particularly in T CD8+ lymphocytes localized in the spleen. Down regulation of this chain is associated with a mechanism of tumor immune evasion in different types of cancer, ^91^ including in ovarian cancer. Here, it is reported that downregulation of that chain is due to a secreted factor of 14 kDa. ^92^ Therefore, increased zeta chain expression in the TCR suggests increased activity of lymphocytes in cancer.

Finally, given the translational nature of the *in vivo* system we used (the humanized mouse model), our data strongly suggests that claudin-4 can be a target of inhibition with CMP to increase the efficacy of the PARPi, niraparib in HGSC patients. Also, the functional link of claudin-4 with SLC1A5/LAT1, particularly LAT1 suggests that targeting claudin-4 could be a way to target indirectly the function of LAT1. Perhaps by modulating the localization of these transporters of amino acids in plasma membrane by stabilizing cell-cell junctions with forskolin97-99

## METHODS

### Cell lines and viral vectors

Human derived cells, OVCA429 (RRID:CVCL_3936), OVCAR3 (RRID:CVCL_DH37), OVCAR8 (RRID:CVCL_1629) were cultured in RPMI-1640 medium (Gibco, Thermo Fisher Scientific, Cat. #11875) plus 10% heat-inactivated fetal bovine serum (Phoenix Scientific, Cat. # PS-100, Lot # 20055-01-01) and 1% penicillin/streptomycin (Corning, Cat. #30-002-CI) at 37°C and 5% CO2. HEK293-FT (RRID:CVCL_6911) were culture similarly but in DMEM medium (Gibco, Thermo Fisher Scientific, Cat. #11995040). pCDH-EF1a-mCherry-EGFP-LC3B (RRID: Addgene_170446); pLenti-Lifeact-tdTomato (RRID: Addgene_64048)

### Inhibition of claudin-4 expression by CRISPRi

Stable transfectant OVCA429 and OVCAR3-dCas9 cells were generated by lipocomplexes (Lipofectamine 2000, ThermoFisher, cat: 11668-019; vector pB-CAGGS-dCas9-KRAB-MeCP2, Add gene: 110824) following the manufacturer’s instructions. Selection was carried out by antibiotic resistance (blasticidin). A gRNA (forward: GCTGGCTTGCGCATCAGGAC; reverse: GTCCTGATGCGCAAGCCAGC) specific for human *CLDN4* (claudin-4) was generated in the Broad Institute portal (https://portals.broadinstitute.org/gppx/crispick/public). Subsequently, the pCRISPRia-v2_base (TagBFP) vector was digested (BstXI and BlpI) to insert the gRNA by ligation, followed by cloning using *E. coli* (Stbl3™, ThermoFisher, cat: C737303), and selection by flow cytometry (MoFlo XDP100; CU Cancer Center Flow Cytometry Shared Resource).

### Lentivirus production and transduction

293FT cells were transfected using lipocomplexes (Lipofectamine 2000, ThermoFisher, cat: 11668-019) containing the viral packaging system of second generation as well as the lentiviral construct of interest pCDH-EF1a-mCherry-EGFP-LC3B (RRID:Addgene_170446), respectively. Supernatant of transfected 293 ft cells was collected, filtered (0.45 µm), used or stored (-80°C).

### Immunoblot and CMP synthesis

To analyze levels of claudin-4 protein expression, tumor cells were scraped from culture plates in the presence of lysis buffer (30 mM Tris HCl pH7.4, 150 mM NaCl, 1% TritonX-100, 10% glycerol, 2 mM EDTA, 0.57 mM PMSF, 1X cOmplete™ Protease Inhibitor Cocktail), placed on a shaker for 10 minutes and spun at 13,000 rpm for 10 minutes. Protein was separated by SDS-PAGE and transferred to PVDF membrane using the TransBlot Turbo (BioRad). Membranes were blocked with Intercept Blocking Buffer (LI-COR, #927-60001) for 2 hour at room temperature. Mouse anti-human claudin-4 (Thermo Fisher Scientific Cat# 32-9400, RRID:AB_2533096, 1:500), rabbit anti-human β-actin (Abcam Cat# ab6276, RRID:AB_2223210,1:10,000), primary antibody incubation was performed overnight at 4 °C. Membranes were washed 3 times for 5 minutes each in TBST (50 mM Tris pH 7.5, 150 mM NaCl, 0.1% Tween-20), followed by secondary antibodies for two hour at room temperature. Membranes were washed again 5 times for 5 minutes each in TBST. For fluorescent detection, bands were visualized using the LI-COR Odyssey Imaging System. CMP was synthesized as previously reported^24^

### Cell cycle analysis by flow cytometry

3x10^5^ cells were seeded onto 6 well-plate, the next day cells were washed (sterile PBS 1x) and changed media. After 24h and 48h incubation cells were detached (trypsin 0.25mM), centrifuged (1800 rpm/5min) and washed with PBS followed by centrifugation. PBS was discarded and cells were fixed using cold ethanol 70% (ethanol, milliQ water, v/v) for 30min (4°C) followed by centrifugation (1200 rpm/5min/4°C). Afterwards, cells were washed twice with PBS and the PBS was discarded after centrifugation (1200 rpm/5min/4°C). Cells were treated with RNAse A (50µL/100µM) for 30min at RT and then stained with propidium iodide 300µL (50µM). Analysis were carried out in the Cancer Center Flow Cytometry Shared Resource, University of Colorado Anschutz Medical Campus.

### ISRE (interferon-stimulated response element) assay

OVCA429 (ISRE; 1x10^5^) cells were seeded on 24 well-plate (1 mL complete RPMI medium). Next day, cells were washed with sterile PBS (1X) and treated with chloroquine, CQ, 40 µM, and 20 µM) and 2’3’-cGAMP (positive regulator of ISRE; InvivoGen, cat: tlrl-nacga23s) in 1mL complete RPMI medium for 24h. The luciferase-luciferin reaction was carried out using the luciferase assay system (Promega, cat: E1501) and bioluminescence was detected by GLOMA MULTI detection system (Promega).

### Immunofluorescence

Cells were fixed with paraformaldehyde at 4% (PBS 1X) for 10 minutes, followed by permeabilization (30 minutes, 0.1% Triton X-100, PBS 1X). Blocking was carried out by 2h incubation with BSA at 5% (PBS 1X, room temperature, RT, shaking). Primary antibodies (mouse, anti-claudin-4, ThermoFisher, cat: 32-9400, at 1:200 dilution; rabbit, anti-GM130, Cell signaling, cat: 12480, at 1:1600 dilution) were incubated (BSA at 2%, PBS 1X) overnight at 4 °C/shaking. Secondary antibodies (AlexaFluor546 anti-mouse, ThermoFisher, cat: A-11030, at 2 µg/mL; AlexaFluor647 anti-rabbit, ThermoFisher, cat: A32733, at 2 µg/mL) were incubated 2h/shaking at RT (BSA at 2%, PBS 1X). Nuclei were stained with DAPI at 1 µg/mL (PBS 1X) for 10 minutes. All microscopy acquisition was carried out in the Neurotechnology Center, University of Colorado Anschutz Medical Campus.

### Generation of HIS-BRGS mice and chimerism evaluation

*Stem Cell Isolation.* HSCs were isolated from PBMCs prepared from clinically-rejected CB units from the University of Colorado Cord Blood bank (Clinimmune Labs, Aurora, CO) using CD34+ magnetic Miltenyi beads, and expanded in short-term cultures with IL6 (10 ng/ml), SCF (40 ng/ml) and FLt3L (20 ng/ml). CD34+ cells, harvested between days 4-6, were frozen in 90% FCS/10%DMSO and stored at −80°C prior to injection into neonate pups. Investigators were blinded from donor identities and the studies were performed in compliance with University of Colorado Institutional Review boards (COMIRB#16-0541).

*HIS-BRGS mice and chimerism evaluation.* To generate human immune system mice, neonatal (d1-3) BRGS (BALB/c*Rag2*^nul*l*^*IL2Rg*^null^*Sirpa*^NOD^) pups, obtained from the laboratory of James Di Santo, were irradiated with 300 rads 2-6 hrs prior to injection with 0.2-0.6x10^6^ expanded then thawed CD34+ cells. The number of cells injected is equivalent to 50,000 fresh CD34+ cells per mouse, i.e. cell count prior to *in vitro* expansion. Mice were bred and engrafted in the University of Colorado Denver Anschutz Medical Campus vivarium with prior Institutional Animal Care and Use Committee (IACUC) protocol and in a facility accredited by the American Association for Accreditation of Laboratory Animal Care. BRGS mice, both breeders and engrafted, were maintained on an alternating bi-weekly Septra-enriched (Uniprim, Harlan) diet. Mice were injected in the facial vein, liver, or both. Humanized mice were generated in the Pre-clinical Human Immune System Mouse model (PHISM) Shared Resource, University of Colorado Anschutz Medical Campus.

All mouse work was performed in accordance with the Guide for the Care and Use of Laboratory Animals and were approved of by the University of Colorado’s Institutional Animal Care and Use Committee (IACUC protocol #283). Humanized Immune System (HIS) mice were i.p. injected with 5x106 cells of a developed patient-derived xenograft (PDX) model, PDX GTFB 1016, described previously. ^68^ Briefly, this model is derived from a primary tumor collected from a patient diagnosed with stage IIIC, who was described to have a high volume of ascites, peritoneal carcinomatosis, and was chemonaïve at the time of sample collection. TP53 and BRCA2 mutations were identified, and we subsequently transfected the cells with a GFP-luciferase tag, enabling tumor tracking by In Vivo Imaging (IVIS, Perkin Elmer), as described previously. ^100,101^ Tumors were allowed to establish for three weeks prior to treatment initiation. After this time, mice were treated with either Niraparib (50 mg/kg, PO) five day per week followed by two consecutive days off or CMP (4 mg/kg, i.p.) every two days. Control mice were treated with respective vehicles on the same treatment schedule (5% DMSO, 30% 300PEG, PO [Niraparib vehicle] and PBS, i.p. [CMP vehicle]). Treatment occurred for 28 days, and mice were IVIS scanned weekly to assess tumor development. One day after the last treatment, mice were euthanized via CO2 inhalation and cervical dislocation, and tissues were collected.

Flow cytometry. Blood, spleen and tumor single cell suspensions were stained, as described previously,^66^ with fluorescently-labeled Abs as listed in Supplemental Table X. Blood samples for chimerism assessment were collected on a Biorad Yeti 5-laser flow cytometer, and spleen and tumor harvest samples were collected on the Cytek Aurora 5-laser spectral cytometer at the University of Colorado Cancer Center Flow Cytometry Shared Resource. All data were analyzed using FlowJo software.

### Statistical Considerations

ImageJ (NIH) and Prism software (v9.0) were used for microscopy and statistical data analysis, respectively. 3 biologically independent experiments were conducted. Unpaired t and Mann–Whitney tests; Kruskal–Wallis test and one-way ANOVA with Dunn’s or Tukey’s multiple comparisons test, respect. Based on normal data distribution, and number of variables. The level of significance was p < 0.05.

#### Abbreviations

HGSC: high-grade serous carcinoma
CMP: claudin mimic peptide
PDX: patient derived xenograft
HIS: human immune system
PARP: Poly (ADP-ribose) polymerase
TIL: tumor infiltrating lymphocytes

## ACKNOWLEDGEMENTS

We acknowledge philanthropic contributions from D. Thomas and Kay L. Dunton Endowed Chair in Ovarian Cancer Research, the McClintock-Addlesperger Family, Karen M. Jennison, Don and Arlene Mohler Johnson Family, Michael Intagliata, Duane and Denise Suess, Mary Normandin, and Donald Engelstad.

## DECLARATIONS

### Ethical Approval

Stem Cell Isolation was performed in compliance with the University of Colorado Institutional Review boards (COMIRB#16-0541). All mouse work was performed in accordance with the Guide for the Care and Use of Laboratory Animals and was approved by the University of Colorado’s Institutional Animal Care and Use Committee (IACUC protocol #283). The University of Colorado has an Institutional Review Board approved protocol (COMIRB#07-935). Reference: ^68^

### Autor’s contribution

BB and MN conceived the overall project. BB supervised the research project. FRV performed the *in vitro* experiments and data analysis along with PW. JL generated humanized mice and performed flow cytometry and analysis. EW and FRV carried out *in vivo* treatments, tumor burden and ascites processing. FRV wrote the manuscript with the contribution of all authors.

### Data availability statement

Data generated in this study are included in this manuscript and in its supplementary information. Datamining and flow cytometry raw data are available upon request to the corresponding author.

## SUPPLEMENTARY FIGURES

**Supplementary Figure 1.**
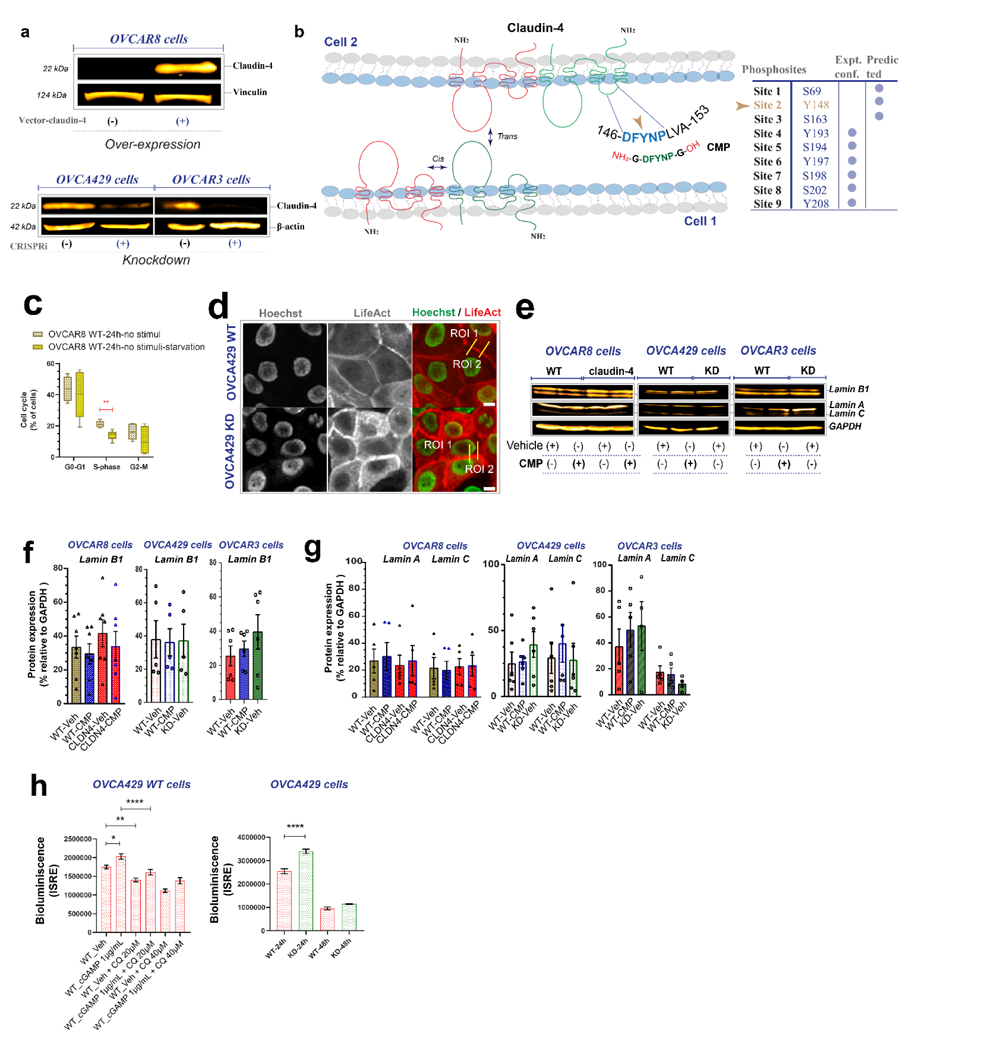
**(a)** Immunoblotting to verify the claudin-4 overexpression (top panel) in OVCAR8 cells (which do not express claudin-4) and its downregulation (knockdown using CRISPRi in OVCA429 and OVCAR3 cells) (bottom panel). **(b)** left, drawing highlighting the structure of claudin-4, in red and green colors (where it can be observed two extracellular loops, large and small) as well as *cis* and *trans* interaction at the plasma membrane as a functional regulation. The drawing also highlights a conserved sequence of claudin-4 (DFYNP) located in the second (small) extracellular loop as well as the sequence of CMP (claudin mimic peptide; DFYNP which is in D-conformation and stabilized by Glycine, thus having NH2 and OH ends flanking the peptide); right, known and predicted phosphosites in claudin-4 (it is highlighted a predicted and uncharacterized phosphosite at Y148, located in the conserved sequence in the small extracellular loop of claudin-4) generated in (http://www.phosphonet.ca/). **(c)** Quantification of cell cycle progression by flow cytometry and propidium iodide-stained OVCAR8 WT cells during normal (RPMI, 10% FBS) culture conditions or starvation (RPMI, 1% FBS). **(d)** Maximum projection of live-cell confocal imaging (xyt) where two different ROIs (region of interest) were used to generate kymographs from junctional F-actin and estimate actin dynamics at multiple time points. **(e)** immunoblotting for lamin B1 and lamin A/C and corresponding quantification relative to loading control (4 independent experiments) in **(f)** and **(g)**, respectively. **(h) Left, t**ransduced OVCA429 WT cells with ISRE vector and treated with cGAPM 1µg/mL (24h) and chloroquine (CQ, 20 and 40µM; 24h) (3 independent experiments; One-way ANOVA and Tukey’s multiple comparisons test; p<0.5); pilot test (one independent experiment using OVAC429 WT and OVCA429 KD-ISRE transduced cells; One-way ANOVA and Tukey’s multiple comparisons test; p<0.5). Graphs show mean and SEM, scale bar 5µm.

**Supplementary Figure 2.**
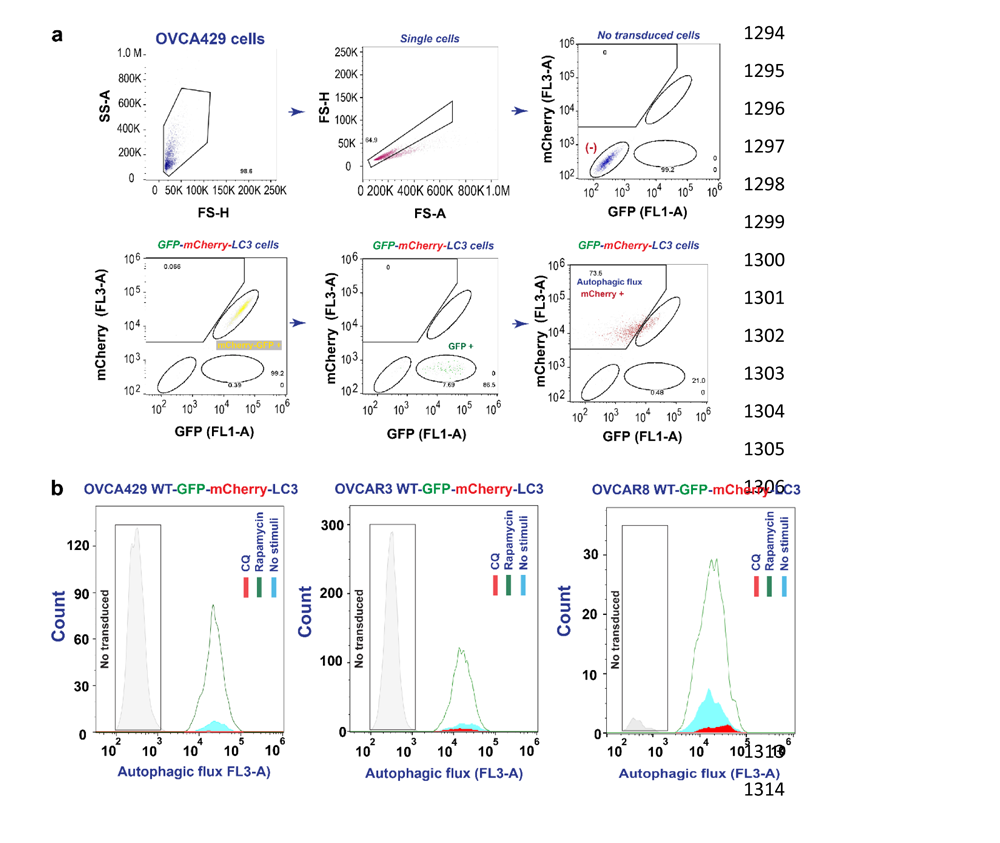
**(a)** Autophagy flux strategy for flow cytometry. **(b)** Confirmation of autophagy flux in HGSC cells (OVCAR8 WT, OVCA429 WT, OVCAR3 WT). CQ, chloroquine (40µM, 24h) was used to show blocking of autophagy; rapamycin, rap (8µM, 24h) was used to show activation of autophagy. Graphs show mean and SEM.

**Supplementary Figure 3.**
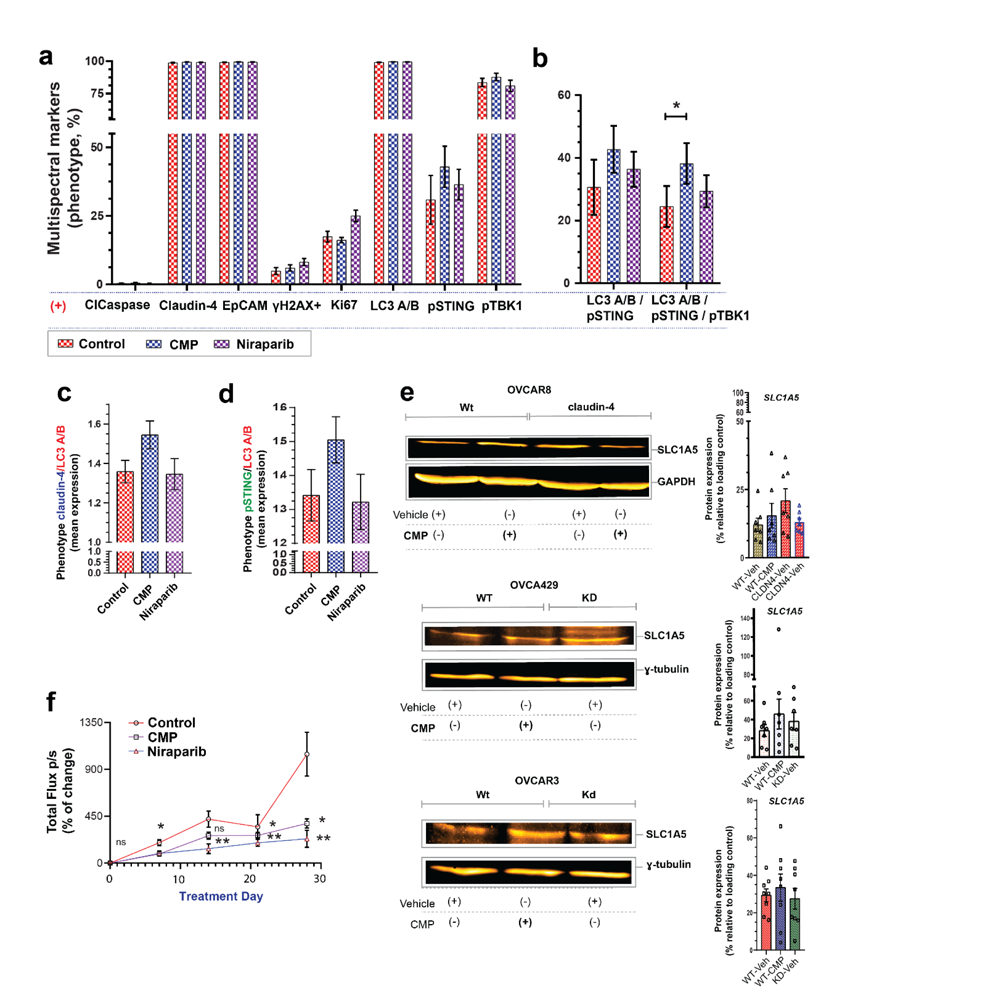
**(a)** Phenotypic frequency (from multispectral immunofluorescence; 8 proteins plus dapi) of cells obtained from ascites generated by HGSC primary tumor in humanized mice from ascites samples. **(b)** Quantification of cells positive for LC3 A/B and pSTING; and LC3 A/B, pSTING plus pTBK1.**(c)** Mean expression of claudin-4 and LC3 A/B and **(d)** pSTING/LC3A/B in ascites samples and (from multispectral immunofluorescence). **(e)** Left, immunoblotting for SLC1A5 in different HGSC cells; right, protein quantification relative to loading control (4 independent experiments). **(f)** Tumor burden (experiment #1) indicated as bioluminescence signal over time. (Two-way ANOVA and Tukey’s multiple comparisons test; p<0.05). Graphs show mean and SEM.

**Supplementary Figure 4.**
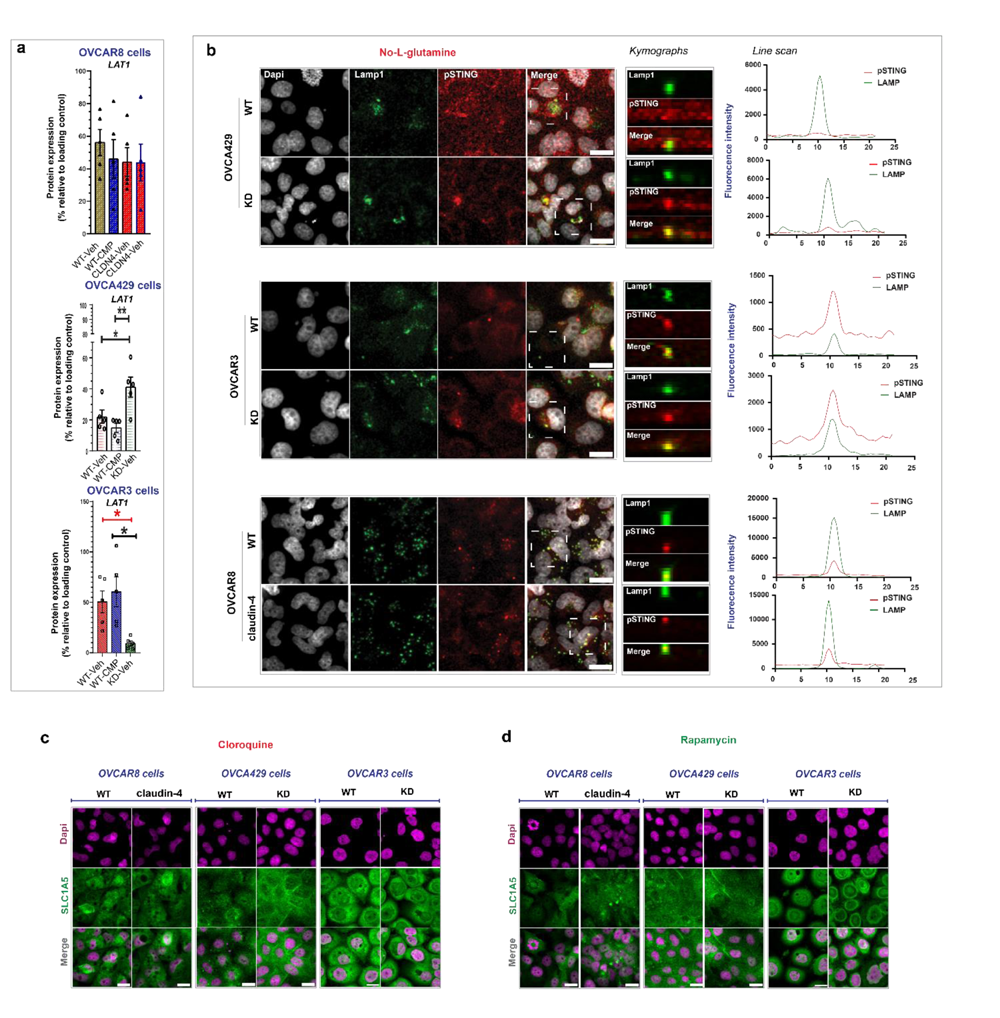
**(a)** Quantification of LAT1 in lysates from HGSC cells relative to loading control (4 independent experiments; One way ANOVA and Tukey’s multiple comparisons test; p<0.05). **(b)** left, confocal images (maximum projections) of HGSC cells cultured without L-glutamine (24h) and stained for pSTING and lamp1; center, kymographs from selected regions (from z-stacks confocal images); right, line scan of selected regions for lamp1 and pSTING. Here is highlighted the co-localization of lamp1 with pSTING before L-glutamine withdrawal. Confocal images (maximum projections) showing SLC1A5 intracellular distribution in HGSC cells before claudin-4 overexpression (OVCAR8-claudin-4) or downregulation (OVCA429 KD and OVCAr3 KD) and autophagy blocking **(c)** (using chloroquine, CQ, 40µM, 24h) or autophagy induction **(d)** (using rapamycin, 8µM, 24h). Graphs show mean and SEM, scale bar 10µm.

**Supplementary Figure 5.**
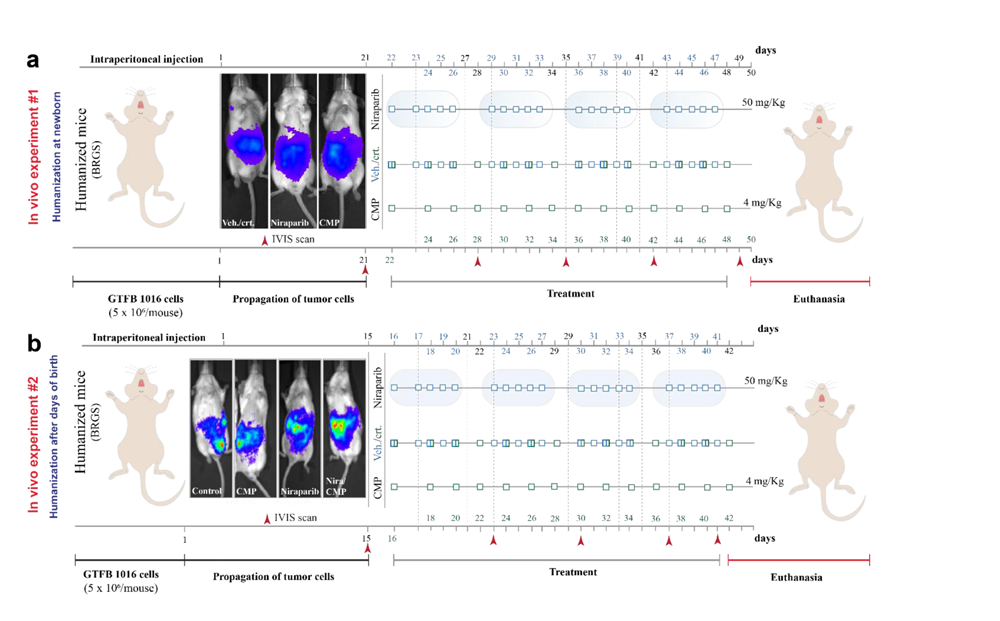
**(a)** and **(b)** Experimental design for targeting claudin-4 in primary tumor in a humanized mouse model (two independent experiments; #1, newborn mice were treated to induced development of human immune system; #2, mice without weaning were treated to induced development of human immune system).

**Supplementary Figure 6.**
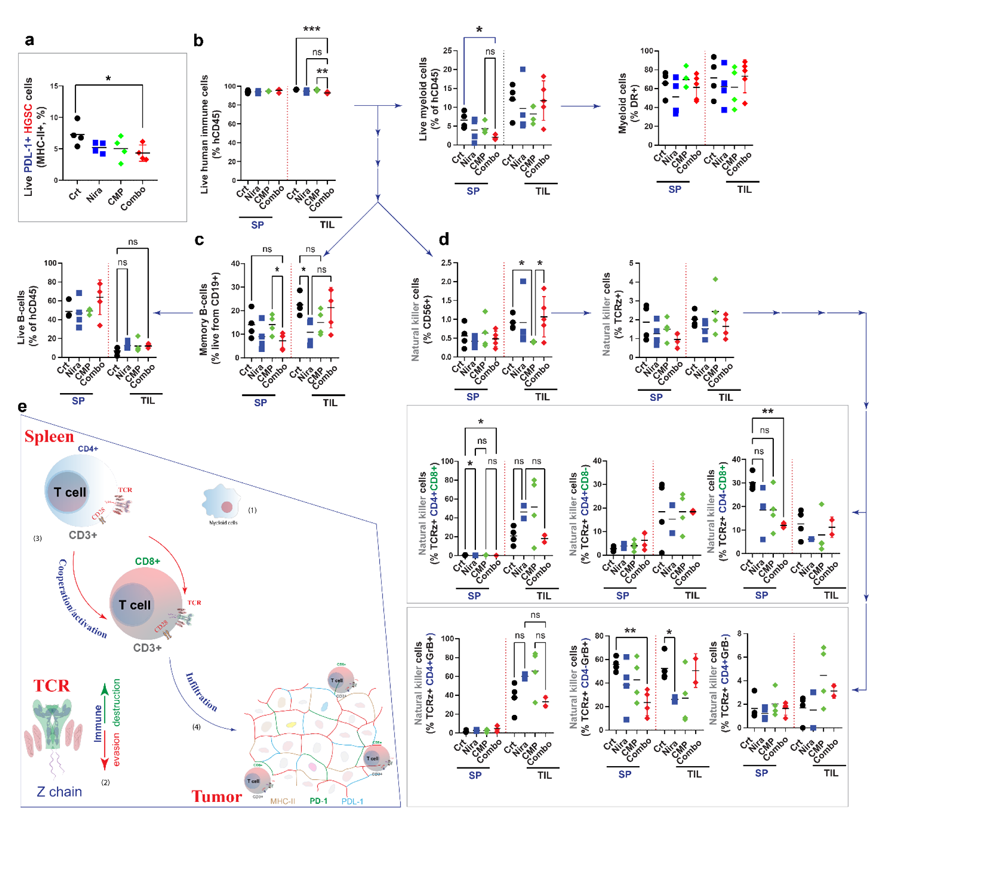
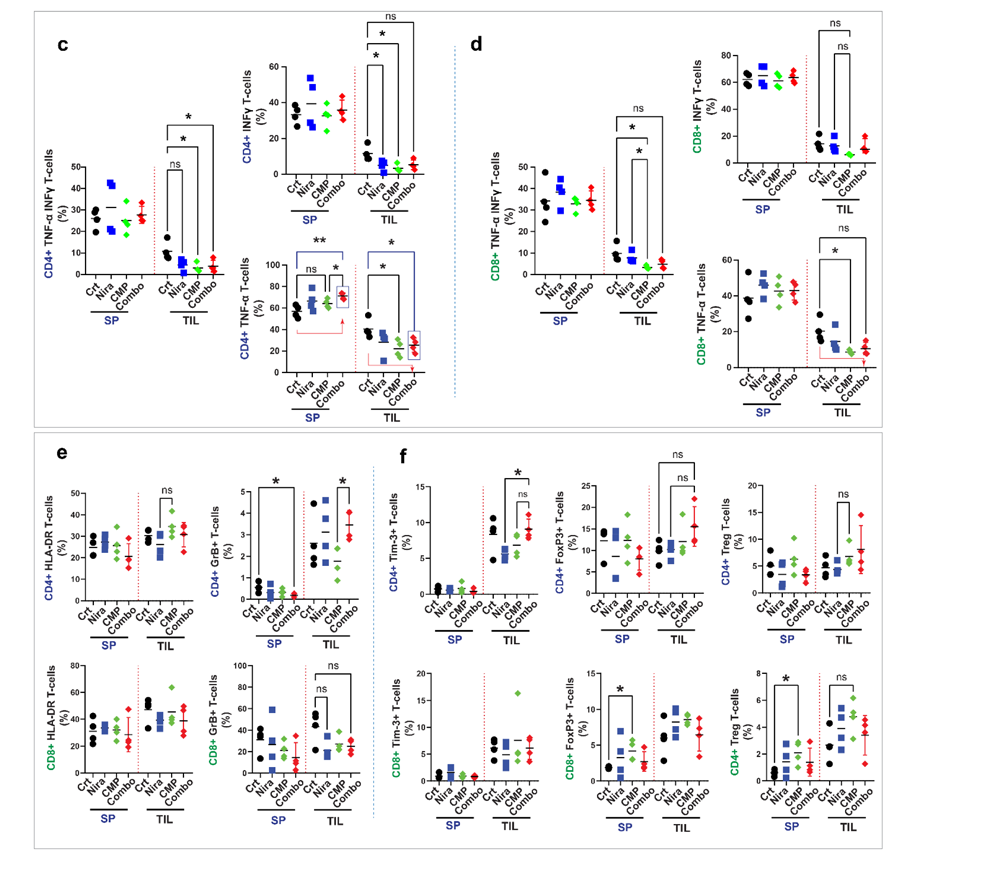
**(a)** Percentage of tumor cells with MHC-II that express PDL-1 expression as well. **(b)** Determination of immune cell populations other than T-cells by flow cytometry. **(c)** and **(d)** Identification of lymphocytes CD4+ and CD8+ producers of TNF-α and INF-ɣ by flow cytometry. **(e)** Proposed model of immune modulation by niraparib and CMP combinatory treatment. Consisting reduced accumulation on myeloid cells in the spleen (1), increased expression of the TCR-zeta chain, which could favor the lymphocyte activation in the spleen as well as the cooperation of T CD4+ and T CD8+ lymphocytes to promote lymphocyte infiltration into the tumor, which show changes in its cellular composition, correlating with reduced tumor growth. () quantification of lymphocytes CD4+ and CD8+ positive for HLA-DR and GrB. **(f)** Identification of lymphocytes CD4+ and CD8+ with expression of Tim3, FoxP3, and Treg (T regulatory cell) phenotype, respectively (All data derives from gate: live human immune cells, T cells, CD4+ and CD8+, respectively). Graphs show mean (every symbol represents data from a different mouse) (n=Control, 4 mice; CMP, 4 mice; niraparib, 4 mice; niraparib/CMP, 5 mice; Brown-Forsythe ANOVA test and Unpaired t with Welch’s correction; p<0.05).

